# Th1 cells are critical tissue organizers of myeloid-rich perivascular activation niches

**DOI:** 10.1101/2024.11.24.625073

**Authors:** Noor Bala, Alexander McGurk, Evan M. Carter, Ikjot Sidhu, Shruti Niak, Scott A. Leddon, Deborah J. Fowell

## Abstract

Aggregating immune cells within perivascular niches (PVN) can regulate tissue immunity in infection, autoimmunity and cancer. How cells are assembled at PVNs and the activation signals imparted within remain unclear. Here, we integrate dynamic time-resolved *in vivo* imaging with a novel spatially-resolved platform for microanatomical interrogation of transcriptome, immune phenotype and inflammatory mediators in skin PVNs. We uncover a complex positive-feedback loop within CXCL10^+^ PVNs that regulates myeloid and Th1 cell positioning for exchange of critical signals for Th1 activation. Th1 cells spend ∼24h in the PVN, receiving initial peripheral activation signals, before redeploying to the inflamed dermal parenchyma. Niche-enriched, CCR2-dependent myeloid cells were critical for Th1 IFNγ-production. In turn, PVN instructional signals enabled Th1s to orchestrate PVN assembly by CXCR2-dependent intra-tissue myeloid cell aggregation. The results reveal a critical tissue organizing role for Th1s, gained rapidly on tissue entry, that could be exploited to boost regional immunity.

**HIGHLIGHTS:**

- Perivascular niche (PVN): myeloid hubs in inflamed mouse and healthy human skin
- Th1 cells enter, get activated, and leave the PVN within first 24h of tissue entry
- Antigen-specific signals in the PVN promote the tissue organizing functions of Th1s
- Th1 cells assemble the PVN via CXCR2-dependent myeloid cell aggregation

## INTRODUCTION

Immunity at barrier surfaces, and other non-lymphoid tissues, requires the inter-tissue recruitment of diverse immune cell types to the inflamed tissue as well as the dynamic intra-tissue clustering of those cells for cooperative exchange of activation signals between neighboring cells^1^. Clustering of immune cells into micro-anatomical niches that nucleate innate and adaptive immune cells is a hallmark of immunity in most inflamed, infected and malignant tissues. Although targeting such microenvironments represents an important goal for enhancing immunity at key anatomical sites, we still do not understand fundamental aspects of niche position, composition and outcome of T cell activation.

Evidence of strategic positioning of immune activation niches has emerged in many tissues including the skin, brain and lung^2–5^. These sites are often at tissue:vascular boundaries^6^, at points of exchange of solutes and cells, serving to expose newly recruited immune cells to regional perturbations in tissue homeostasis from infection or tissue damage. They can be present in the steady state such as the adventitial cuff of the lung and liver^7,8^ and the dural sinus of the brain^9,10^, but can also be induced upon immune challenge like the de novo generated perivascular niches (PVN) in the skin^11,12^. Perivascular niches are often adjacent to postcapillary venules where leukocytes are thought to enter inflamed tissues^5,13^. Common to many of these regional immune niches is the aggregation of myeloid cells, expression of a variety of chemokines and the co-clustering of effector T cells.

Recent studies highlight the disparate fates of T cells activated in PVNs, from boosting of effector functions like IFNγ production from Th1 cells in cutaneous and neuro-inflammation^5,11,12,14–16^, to supporting stem-like T cells in cancer^17^ that may either improve^18,19^ or exclude/suppress anti-tumor immunity^20,21^, to harboring stem-like CD4 T cells in autoimmunity that contribute to chronicity^22^. These associations between induction of PVNs and the modulation of T cell function suggest this regional control point could be an important therapeutic target for regulating tissue immunity, yet we lack a basic understanding of the spatial and temporal dynamics of T cell activation in these niches and the cellular players involved in their intra-tissue assembly.

Single cell transcriptomic data has revealed an incredible degree of heterogeneity within and between individual immune cell types in inflamed tissues. Mechanistic understanding of such heterogeneity will require the ability to determine how cell interactions are shaped by spatial location and by time in tissue^17,23–25^. To determine the impact of time and space on the early events of initial T cell activation following recruitment to the inflamed skin, we integrated dynamic time-resolved IV-MPM imaging with a new multiphoton-guided platform for spatially-resolved single cell transcriptomics (Niche-seq), multiparameter flow cytometry (Niche-flow) and immuno-protein profiling (Niche-secretome). This approach uncovered a complex exchange of signals within the perivascular space that regulates myeloid and T cell positioning to optimize early T cell activation at the point of upon tissue entry. We establish the CXCL10^+^ perivascular niche as an initial site of antigen-specific Th1 activation, within 24h of tissue entry, before Th1 cells move into the dermal tissue parenchyma. The PVN is enriched in myeloid cells that share a common niche-specific inflammatory program that underpins a chemokine-enriched hub for immune cell recruitment and activation. The myeloid PVN transcriptional signature was sufficient to identify similar perivascular niches in healthy human skin highlighting the importance of these immune aggregates in immunosurveillance across species. Th1 activation in the PVN was CCR2-dependent. Instructional PVN signals led to an unexpected Th1 gain of function, driving the upregulation of myeloid-recruiting chemokines in Th1 cells specifically at the niche, suggesting the T cells themselves may play a role in the construction of the PVN. Indeed, Th1 cells were necessary and sufficient for robust PVN assembly in a CXCR2-dependent manner. That effector T cells can assemble their own activation niches at inflamed sites, through the intra-tissue repositioning of activated myeloid cells, suggests this new process could be exploited to boost local immunity in chronic infection and cancer, or be therapeutically disabled in the setting of immune pathology and autoimmunity.

## RESULTS

### CXCL10-expressing perivascular niches serve as the first step in peripheral Th1 antigen-specific activation

Induction of IFNγ-mediated Type 1 dermal inflammation following immunization (protein in Complete Freund’s Adjuvant (CFA)^12^, microbial challenge (*Leishmania major, Staphylococcus aureus*)^12,16^ or contact hypersensitivity^11^ induces the formation of post-capillary PVNs, that we previously defined by the clustering of CXCL10^+^MHC-II^+^ myeloid cells^12^ (Figure 1A). These dermal PVNs appear to optimize CXCR3^+^ Th1 cell antigen encounter upon entering the inflamed tissue (Figure 1A), but we are lacking a temporal understanding of how long Th1 cells spend in the niche on tissue entry and the functional consequences of initial Th1 activation within these perivascular niches.

**Figure 1.**
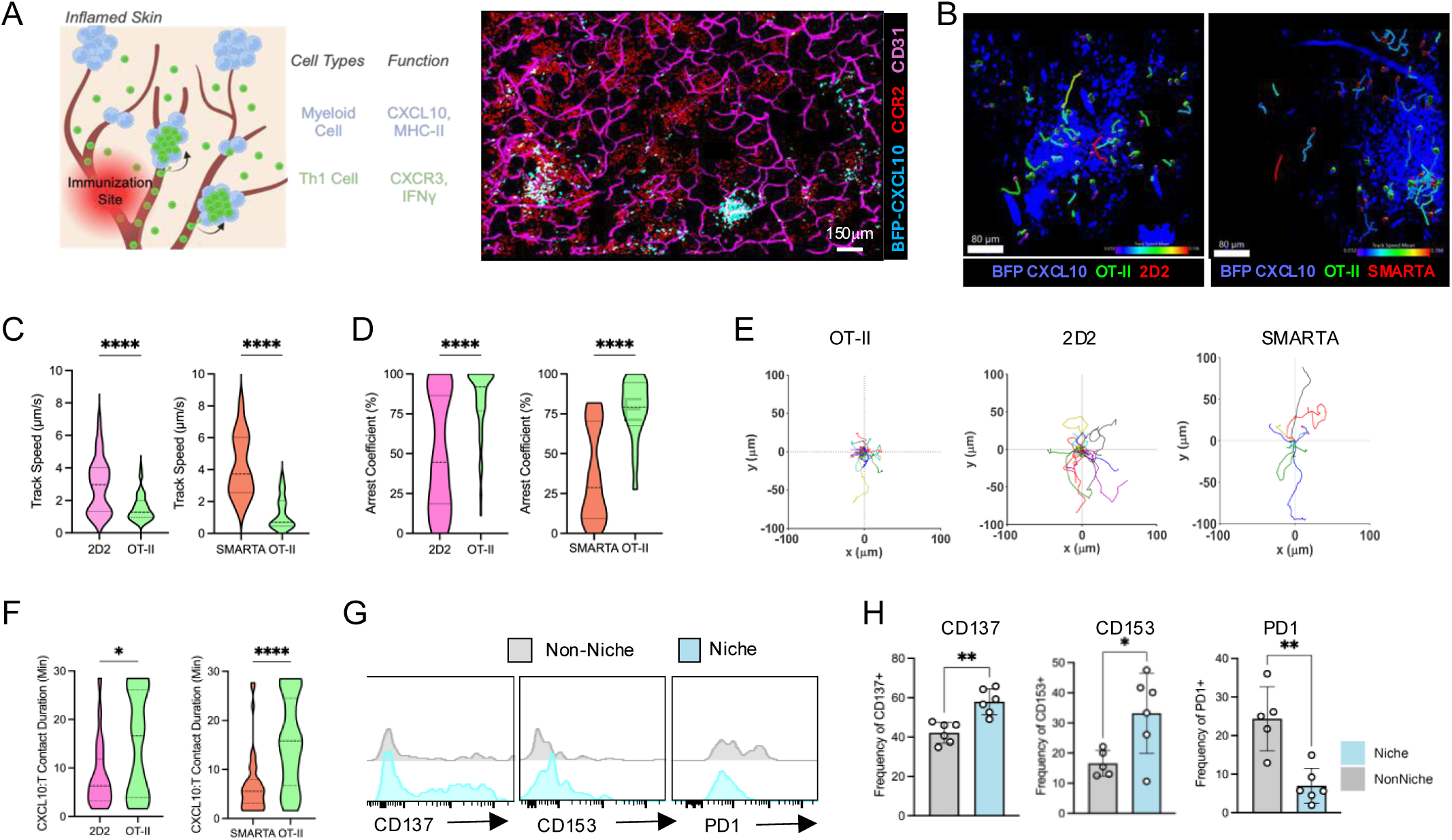
Antigen Specific activation of Th1 cells in the CXCL10^+^ perivascular niche. (A) Schematic of dermal perivascular niches (PVN) in the inflamed dermis (left) and IV-MPM image (right) of the distribution of CXCL10^+^ PVNs in the dermis of REX CXCL10-BFP X CCR2-RFP^+/-^ mice following OVC/CFA immunization. (B-F) 2.5 x 10^6^ OT-II and 2D2 or OT-II and SMARTA Th1 cells were co-transferred i.v. into OVA/CFA-immunized REX3 reporter mice, and their dynamic movement and position relative to the CXCL10^+^ PVNs determined on d5 of immunization by IV-MPM of the inflamed ear dermis. (B) Images of distribution of antigen and non-antigen specific Th1 cells with respect to PVN. Scale bar, 80 µm. Representative data from three independent experiments, > 8 imaging volumes; >30 tracks per plot. (C and D), Track mean speed (C) and arrest coefficient (D) of OT-II and 2D2 or SMARTA Th1 cells inside of the PVNs. (E) Migratory paths (x-y projections) of OT-II, 2D2, and SMARTA Th1 cells with respect to CXCL10^+^ PVN. (F) CXCL10^+^ cell:T-cell contact time according to antigen specificity, 30 min IV-MPM. (G and H) Representative histograms (G) and frequency (H) of CD137^+^, CD153^+^, and PD1^+^ Th1s in Non-Niche and Niche regions of the OVA/CFA-immunized dermis. PVNs were identified in the inflamed dermis on d5 p.i. via IV-MPM, mapped, and biopsied for ex vivo analysis via flow cytometry. Gated on CD4^+^/Live/Lymphocytes/CD45.1^+^. Statistical analysis performed using Mann-Whitney test; *p<0.05, **p<0.01, ***p<0.001, ****p<0.0001.

To determine Th1 dynamics and duration of antigen-specific events within the PVN, we utilized a co-adoptive transfer model into immunized CXCL9/10 reporter mice (REX3)^26^ and intra-vital multiphoton microscope (IV-MPM) imaging. *In vitro* generated MOG-specific (2D2)^27^ or LCMV GP_61-80_-specific (SMARTA)^28^ Th1 cells were intravenously co-transferred alongside ovalbumin-specific (OT-II) Th1 cells into REX3 mice intradermally immunized with OVA/CFA in the ear pinna. Day 5 post-immunization (p.i.) IV-MPM was used to visualize the inflamed dermis and identify the CXCL10-BFP^+^ PVN (Figure 1B). Non-antigen specific (2D2 & SMARTA) Th1 cells entered the inflamed dermis but displayed increased speed (track speed mean) at the niche compared to their antigen-specific OT-II counterparts (Figure 1C, Video S1), suggesting the presence of antigen at the PVN reduces Th1 migration and promotes Th1:APC interactions. Consistent with that notion, OT-II Th1s had a higher arrest coefficient at the niche compared to 2D2 and SMARTA Th1 cells (Figure 1D, E) and prolonged contact times with CXCL10-BFP^+^ cells in the PVN (Figure 1F). To further examine spatially-defined activation events in the inflamed skin, we focused on antigen-specific OT-II cells and developed an IV-MPM-directed system to isolate biopsies of niche-enriched and non-niche tissue areas (see workflow in Figure 3, Figure S1) for ex-vivo flow cytometry. Co-stimulatory activation markers from the TNF super family, CD137 (4-1BB) and CD153 (CD30L), were found to be increased in niche-specific Th1 cells whereas the inhibitory receptor PD-1 was low on PVN Th1s but higher on non-niche Th1 cells (Figure 1G, 1H). Previous reports have shown that increased PD-1 expression on OT-II CD4 T-cells is associated with antigen disengagement and increased motility in the inflamed dermis^29^, therefore our data are consistent with the PVN being a site of early Th1 cognate activation prior to further tissue exploration.

To establish the spatiotemporal dynamics of Th1 cells in the PVN, we first used Th1 cells expressing the photoconvertible Kaede protein^30^ for time-resolved assessment of location and activation status (Figure 2A). Kaede-green OT-II Th1 cells were transferred to mice and immunized with OVA/CFA, on d5 Th1 cells recruited to the ear were photoconverted to Kaede-red by exposure of the ear pinna to 405 nm light, flow cytometry was used to confirm conversion of all Th1 cells to Kaede-red post-conversion (Figure 2A).). After 24h, the previously present Kaede-red Th1 cells could readily be distinguished from the Kaede-green Th1 cells that had entered the inflamed ear since photoconversion (Figure 2A). To determine spatial-distribution with time in tissue, the inflamed dermis was visualized using IV-MPM to first determine location of the converted Kaede-red Th1 cells immediately post-photoconversion (Figure 2B). Re-examining the same tissue 24h later, we observed that most of the previously recruited Kaede-red Th1 cells were no longer localized to the niche, while newly recruited Kaede-green Th1 cells were mostly localized to the PVN (Figure 2B, quantitation right panels). Consistent with the niche-flow data (Figure 1G, 1H), Kaede-green

**Figure 2.**
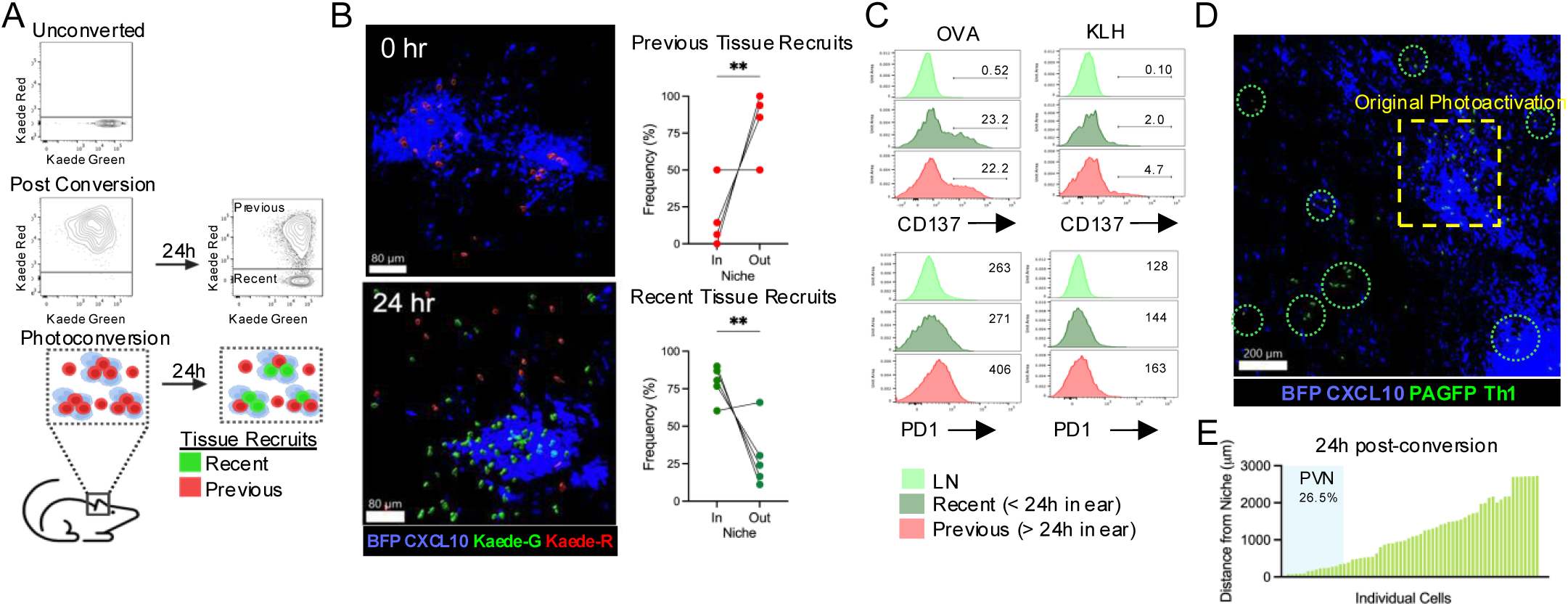
Time-resolved Th1 entry and exit of the PVN in the first 24h from tissue entry. (A) Schematic of Kaede Th1 time-stamp model. OT-II^+^ Kaede^+^ Th1 cells were transferred into OVA/CFA-immunized REX3 mice. On d5, p.i., all Th1 cells in the inflamed ear were photoconverted to Kaede-red on exposure to 405 nm light. 24 h later, the ear tissue contained Recent recruits, Kaede-green Th1 cells (< 24h in ear), and Previous tissue recruits, Kaede-red Th1 cells (> 24h in ear). (B) IV-MPM images (left) of the distribution of OT-II^+^ Th1 cells relative to the PVN immediately post-photoconversion (0h) or 24h after photoconversion (24h): Kaede-green, Recent recruits; Kaede-red, Previous recruits. Right, quantitation of the frequency of Kaede-green and Kaede-red within and outside of the PVN, 24h after photoconversion. Representative data from three independent experiments. Statistics by Mann-Whitney, **p<0.01. (C) Kaede OT-II Th1 cells were transferred into WT albino animals immunized with either OVA/CFA or KLH/CFA. The inflamed ear was photoconverted 24h prior to harvest on d6 p.i. Representative histograms of frequency (CD137) or MFI (PD-1) by OT-II Th1 cells prior to tissue entry (lymph node, LN), 24h post tissue entry (Recent, Kaede-green) and after persisting within the inflamed dermis for >24h (Previous, Kaede-red), with (OVA) or without (KLH) cognate antigen. Representative data from four independent experiments. (D-E) Spatiotemporal tracking of PVN-specific Th1 cells. OT-II PA-GFP^+^ Th1 cells were transferred into OVA/CFA-immunized mice. On d5, single CXCL10^+^ PVNs were identified per mouse using IV-MPM and 830nm laser applied to activate GFP within the PVN localized Th1s. 24 h later, the original photoactivation site was reidentified, example image (D), and distances of PA-GFP^+^ Th1 cells from the PVN calculated (E). Representative data from three independent experiments.

OTII Th1 cells that had recently entered the skin (<24h in ear) rapidly expressed the activation marker CD137 if their cognate antigen, OVA, was present (OVA/CFA vs. KLH/CFA). The photoconverted Kaede-red cells previously residing in the tissue (>24h in ear) maintained CD137 but also expressed PD-1 (Figure 2C). These data suggest Th1 cells initially accumulate in the PVN on tissue entry, are activated, and then reposition outside of the PVN within 24h. To directly map Th1 re-location with time in the tissue, we utilized OT-II Th1 cells expressing photoactivatable GFP (PA-GFP)^31^, enabling us to mark Th1 microanatomical location to the PVN and follow subsequent redistribution. PA-GFP OT-II Th1 cells were adoptively transferred into immunized REX3 mice. For each mouse, the PVN was visualized using IV-MPM and corresponding Th1 cells in the PVN precisely photoactivated by exposure to 830 nm irradiation with the MPM laser. Hair follicles were used to register the PVN tissue location allowing us to revisit the same niche 24h later. While some Th1 cells remained at the niche, the majority of Th1 cells originating in the PVN had migrated into the tissue parenchyma within 24h, often >1000 microns from the PVN (Figure 2D, 2E). Taken together, these dynamic spatiotemporal studies establish the PVN as a site of initial antigen-specific Th1 activation in the inflamed dermis, within 24h of tissue entry, followed by movement deeper into the tissue parenchyma.

#### System-level characterization of the PVN identifies a chemokine-rich myeloid hub

To determine the exchange of signals within the PVN that shape initial Th1 activation, we developed a new systems-level approach to understand the biological complexity of immune activation *in situ* in the PVN. We integrated IV-MPM single cell dynamics with spatially resolved single cell transcriptomics, flow cytometry and immuno-protein profiling to identify key regulators in the choreography and functional programming of lymphocytes specifically in the PVN (Figure 3A). REX3 mice were immunized with CFA/OVA and OT-II Th1 cells adoptively transferred, on d5 post-immunization we imaged the tissue by IV-MPM for CXCL10^+^ aggregates located perivascularly, based on acute CD31 staining immediately prior to imaging. The spatial coordinates for CXCL10^+^ PVN (niche) and non-niche regions of the inflamed ear pinna were mapped using the microscope user interface (Figure S1A). We optimized the size of the biopsy to ensure full niche capture from the imaging coordinates, with a 2mm punch biopsy providing sufficient tissue margins around the PVN for accurate niche isolation, and often capturing 2-3 adjacent niches in the same biopsy (Figure S1A, B). Non-niche regions were identified in a similar fashion, ensuring the absence of CXCL10^+^ PVN aggregates (Figure S1). Niche-enriched and non-niche region biopsies were analyzed for spatially-restricted differences using single-cell RNA sequencing (Niche-Seq), flow cytometry (Niche-Flow), and Luminex cytokine/chemokine multiplexed bead array (Niche-Secretome).

**Figure 3:**
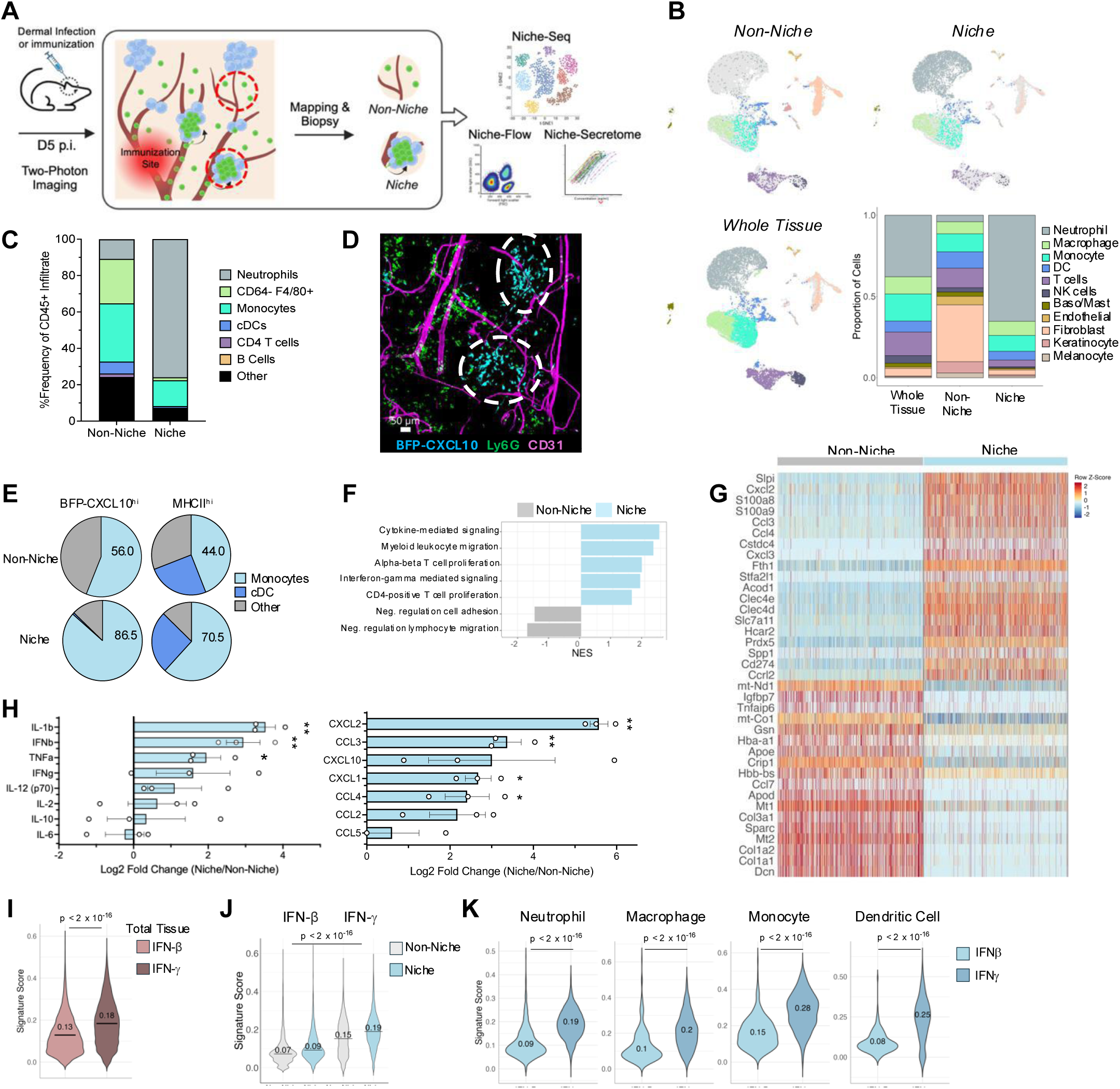
Spatially-resolved PVN cellular composition and inflammatory milieu. (A) Schematic of integrated in vivo and ex vivo analysis workflow for characterization of the PVN. OT-II Th1 cells are transferred into OVA/CFA-immunized REX3 mice and the inflamed dermis imaged on d5 by IV-MPM. Niche and Non-Niche regions are mapped on the MPM user interface and resulting coordinates are used to biopsy the respective regions for ex-vivo analysis in the form of scRNA-seq (Niche-Seq), flow cytometry (Niche-Flow), and Luminex bead assay for inflammatory cytokines and chemokines (Niche-Secretome). (B) Integrated uniform manifold approximation and projection (UMAP) of identified cell types within the Niche-Seq data set and their distribution within whole tissue, non-niche, and niche regions (stacked bar graph). Representative data from two independent experiments: n = 2 whole tissue; n = 6 (pooled) niche; n = 6 (pooled) non-niche; n = 19,737 cells. (C) Compositional analysis of immune cell types in Non-Niche and Niche regions by flow cytometry, Niche-Flow. Representative data from three independent experiments. (D) Representative IV-MPM image of distribution of neutrophils (green, FITC-anti-Ly6G mAb), CXCL10-BFP^+^ cells (blue) and CD31 vessels (magenta, AF647-anti-CD31 mAb) in the d5 OVA/CFA inflamed dermis of REX3 mice; PVN highlighted with a white dotted circle. (E) Niche-Flow on d5 OVA/CFA inflamed dermis of REX3 mice, gated on CXCL10-BFP^hi^ and MHC-II^hi^ cells. Data shown is representative of three independent experiments. (F) Gene Set Enrichment Analysis (GSEA) on the scRNAseq data for Niche and Non-Niche regions. GSEA was performed on ranked gene list with p-value cutoff 0.05; normalized enrichment score (NES). (G) Single cell top 20 differentially enriched genes within the Niche and Non-Niche regions. Differential expression was calculated and filtered adjusted p value, *p<0.05. (H) Niche-secretome, Luminex multiplexed bead-based immunoassay of inflammatory mediators in Niche and Non-Niche biopsy lysates from the d5 OVA/CFA inflamed dermis of REX3 mice. Log2 fold change Niche to Non-Niche lysates from three independent experiments. Statistical analysis performed using multiple unpaired t-test; *p<0.05, **p<0.01. (I-K) Signature scores for responsiveness to IFN-β or IFNγ cytokines: (I) whole tissue scRNAseq data; (J) Niche and Non-Niche regions; (K) myeloid cell clusters in niche region.

For the single cell Niche-seq, we collected niche-enriched and non-niche regions as well as whole tissue (the inflamed ear pinna). Dimension reduction, unsupervised clustering and post hoc annotation on the integrated data set identified 11 distinct cell types based on expression markers from pangloaDB^32^ as well as previous reports that have characterized the inflamed skin^33^ (Figure S2A-C, Figure 3B). Of the integrated data set, neutrophils (∼35%) were the dominant immune cell type, followed by macrophages (∼13%) and monocytes (∼14%), with T cells and NK cells representing ∼14% and ∼10% respectively. Dendritic cells made up just 6% of the cellularity. Non-lymphoid cells, endothelium, fibroblasts, keratinocytes and melanocyte were all detected but at low frequencies (0.2-4.7%) (Figure S1C). The identified cell types were preferentially located in either niche-enriched or non-niche regions, with the niche primarily a hub for myeloid cells (∼85% neutrophils, macrophages and monocytes). The myeloid-rich PVN cellular composition was confirmed by Niche-flow (Figure 3C). The typical niche-enriched biopsy contains some tissue margins to ensure accurate PVN capture (Figure S1), therefore we used MPM to assess the spatial positioning within niche regions of the main myeloid cell subsets identified by Niche-seq and Niche-flow. Neutrophils and monocytes occupied distinct spatial regions: CXCL10^+^ monocytes defining the PVN and neutrophils positioned ‘niche-adjacent’ on the margins of the PVN (Figure 3D). Niche-flow revealed a striking accumulation of CXCL10^hi^ and MHC-II^hi^ inflammatory monocytes (CD64^+^F4/80^+^CD11c^+^ or CD11c^−^) in the PVN compared to classic dendritic cells (cDC), mainly cDC2s (CD64^−^F4/80^−^CD11c^+^CD26^+^XCR1^−^)^12^ (Figure 3E). Thus, the PVN is enriched in monocyte-derived APCs that could activate incoming effector T cells. Consistent with previous reports of inducible skin associated lymphoid tissues (iSALT)^34^, the perivascular niches did not contain appreciable numbers of B cells.

GSEA of genes enriched in the niche identified biological processes associated with myeloid recruitment and T cell activation (Figure 3F). Indeed, examination of the top 20 genes within each region (non-niche vs. niche) revealed upregulation of many chemotactic factors in the niche related to the recruitment and positioning of macrophages, monocytes, and neutrophils (Ccl3, Cxcl2, Cxcl3, Ccl4, Ccrl2) (Figure 3G). Other genes enriched in the niche included genes associated with innate immunity (Hcar2, Clec4e, Clec4d, Slc7a11, Acod1) and immune regulators such as the IL1R antagonist, Il1rn, and endopeptidase inhibitors (Slpi, Cstdc4, Stfa2l1). To better understand the spatial compartmentalization of inflammatory mediators in the dermis, we isolated tissue extracts from niche and non-niche regions for immuno-protein profiling by Luminex (ThermoFisher). Consistent with the scRNAseq data, the PVN contained elevated levels of proinflammatory cytokines IL-1β, IFNβ, TNFα and IFNγ and chemokines for myeloid cell recruitment, CXCL2, CCL3, CCL4 and CXCL1 (Figure 3H). The presence of both innate and adaptive inflammatory mediators in the PVN, prompted interrogation of the differential responsiveness to these cytokines in the niche by applying type 1 (IFNβ) and type II (IFNγ) interferon-responsive gene signatures^35^ to the Niche-seq data set (Figure 3I-K). IFNγ-responsiveness dominated the inflamed tissue (Figure 3I, Figure S2D), although both IFNγ and IFNβ gene signatures were enriched in the niche relative to non-niche regions (Figure 3J). The monocytes and DCs had the highest signature score for IFNγ genes in the niche (Figure 3K) and, consistent with the elevated expression of MHC-II and CXCL10 in the PVN (Figure 3E), those monocytes expressing high levels of CXCL10 had a higher IFNγ signature score (Figure S2E). Thus, the PVN is a hub for expression of interferon-γ-induced chemokines and cytokines that can enhance the microanatomical clustering of myeloid cells, nucleating the accumulation of monocytes with the capacity to activate effector T cells.

### A cooperative myeloid activation program in the PVN

To determine the potential for myeloid cooperativity in the PVN, we analyzed the scRNAseq data for connectivity and shared functional programs. Each myeloid cell type in the niche exhibited a distinct pattern of upregulated chemokines, revealing both common and specialized recruiting potential (Figure 4A). All cell types contribute to the Cxcl3 enrichment in the niche, while Cxcl2 niche enrichment came from macrophage, monocyte and dendritic cell sources. Neutrophils, macrophages and dendritic cells, but not monocytes, were enriched for Ccl3 and Ccl4 gene expression in the PVN, while monocytes remained the key source of Cxcl10 as confirmed by our niche-flow (Figure 3E). We utilized CellChat^36^ to infer niche-specific cellular communication networks based on ligand and receptor expression (Figure 4B, C). By collectively analyzing the Ccl and Cxcl pathways, we identified predicted “Senders” and “Receivers” in each region. In the non-niche region, fibroblasts were the primary “Senders,” while neutrophils, macrophages, and monocytes were the primary “Receivers.” Conversely, within the niche region, neutrophils, macrophages, and monocytes were the primary “Senders,” and T-cells were the primary “Receivers” (Figure 4B). For Ccl chemokines, niche-enhanced interactions were primarily between neutrophils and macrophages, neutrophils and monocytes, and autocrine signaling within neutrophils (Figure 4C). With neutrophils appearing to dominate communication in the PVN, despite being positioned on the PVN margins (Figure 3D), we tested whether neutrophils were required for niche formation and/or maintenance. Using an enhanced neutrophil-depletion strategy^37^, we were able to significantly reduce neutrophils in the circulation and in the inflamed dermis (Figure S3). However, neutrophil depletion had no effect on the ability of monocytes to be recruited to the inflamed dermis, nor in their ability to cluster into CXCL10^+^ PVNs (Figure S3). In contrast, in the Cxcl pathway, niche interactions were primarily between T-cells, monocytes, macrophages and DCs (Figure 4C).

**Figure 4:**
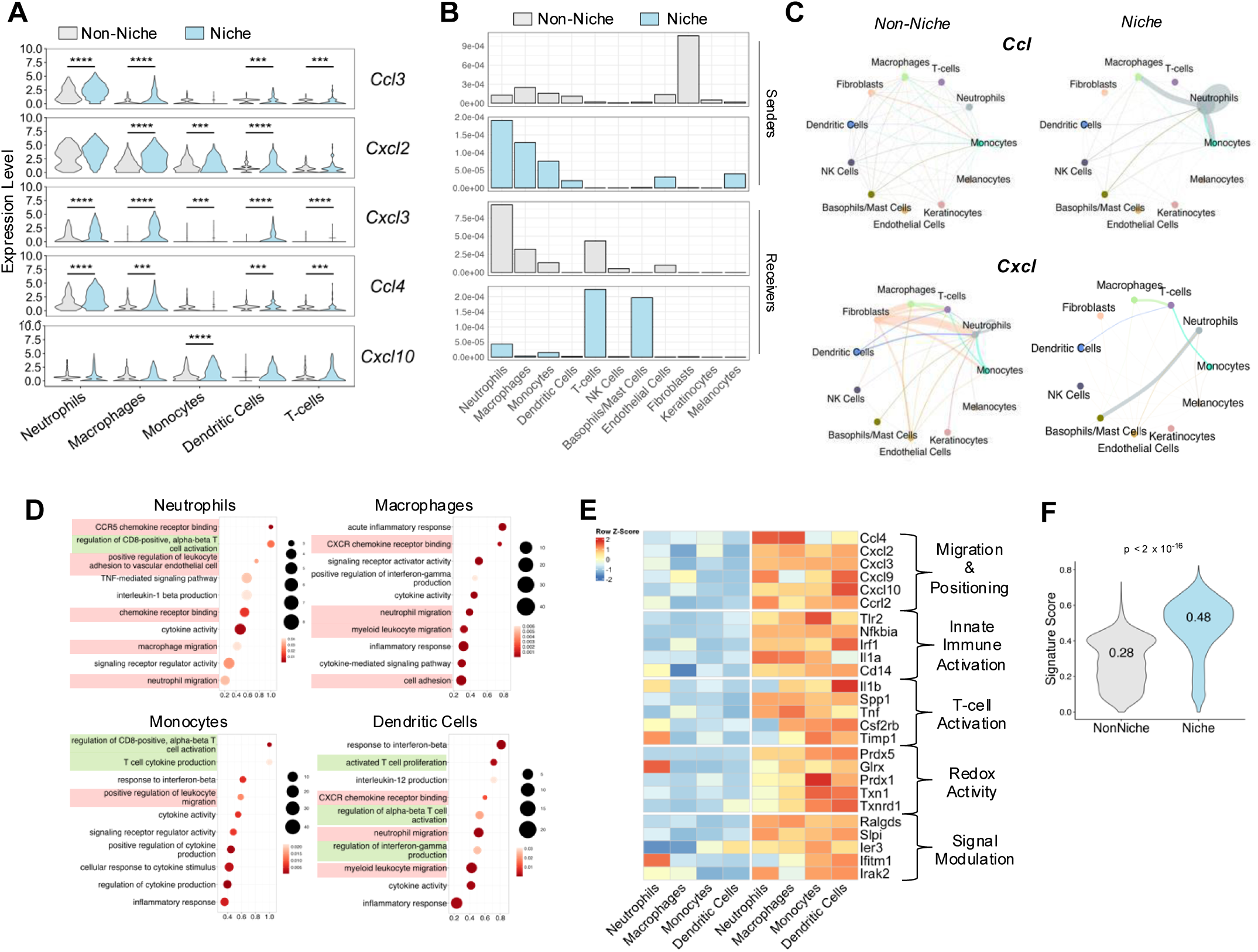
Cooperative clustering of myeloid cells with common activation programs at the CXCL10^+^ PVN. scRNAseq data analysis from d5 OVA/CFA dermis. (A) Expression levels of top chemotactic genes by myeloid cell subsets within Niche and Non-Niche regions. Statistics by Wilcoxon Rank Sum test; ***p<0.001, ****p<0.0001. (B) Communication probabilities of CCL and CXCL pathways for ligand-expressing (Sender) and receptor-expressing (Receiver) cell clusters in Niche and Non-Niche environments using CellChat. (C) Circle plots depicting the differential number of interactions between identified cell clusters in Niche and Non-Niche environments for ligand-receptor pairs within the CCL and CXCL pathways using CellChat. Line thickness represents interaction strength. (D) GSEA analysis for neutrophil, macrophage, monocytes, and dendritic cell clusters, performed on ranked gene lists from differential expression analysis of Niche versus Non-Niche, adjusted p-value, p<0.05. (E) Gene signature derived from the leading edge of GSEA results across myeloid cell clusters, includes genes overlapping in the leading edges of at least 3 out of 4 myeloid cell subsets. (F) Signature score from (E) applied to Niche and Non-Niche scRNAseq data. Statistical analysis performed using a pairwise T-test comparing group levels.

To understand the functional status of cells in the niche we performed gene set enrichment analysis (GSEA) on the individual myeloid cell clusters. The results indicated that cells within the niche were enriched for pathways involved in leukocyte migration/recruitment, T-cell activation, and cytokine responses, with many terms commonly enriched across the cell types (Figure 4D). Genes enriched in the PVN in at least 3 out of the 4 myeloid cell types were identified and used to define a PVN gene signature (Figure 4E) that contains the enriched chemokines, myeloid and T cell activation genes, including those involved in pattern recognition, and regulators of redox activity. Applying this myeloid signature back to niche and non-niche scRNAseq data sets confirmed enrichment of the gene signature in the PVN (Figure 4F). These gene expression patterns in the PVN are consistent with local recruitment and activation of pro-inflammatory myeloid cells for Th1 effector T cell activation.

Unlike mice kept under SPF conditions, human skin has higher commensal microbe colonization as well as experiencing frequent exposure to potential pathogens, hence is a more immunologically active tissue in healthy individuals. Indeed, clusters of DCs, T cells and macrophages forming a perivascular sheath have been described in healthy human skin using 3D whole mount immunofluorescence staining techniques^38^, and were predicted to mark microdomains where immune interactions can occur. More recent 3D reconstruction tools have been applied to serial sections of human skin and has enabled quantitation of T cell location, with most T cells found within 25μm of the nearest endothelial cell^39^. However, the functional status of the vascular-associated immune aggregates remains unresolved. To determine if the induced PVN myeloid activation program identified in our mouse model could identify similar activation niches in healthy human skin, we applied the PVN myeloid gene signature (Figure S4) to spatial transcriptomics (ST) data from healthy human skin biopsies^40^. Healthy skin spot transcriptome clustering was concordant with distinct anatomical neighborhoods in the skin (Figure 5A, left), with T cells and myeloid cells enriched in immune niches in the perifollicular, hair follicle and dermal vasculature clusters as previously described^40^. We applied the PVN myeloid gene signature to the skin ST clusters and visualized the distribution by UMAP (Figure 5A, right) and determined the signature score across ST clusters (Figure 5B). The PVN myeloid gene signature was most enriched in the perifollicular and dermal vasculature clusters (Figure 5B, 5C). Thus, the PVN myeloid gene signature identified from inflammation-induced PVNs of the mouse skin can identify similar spatially-restricted perivascular myeloid cell activation in healthy human skin. These data highlight conserved perivascular myeloid activation niches in mouse and human skin, and identify similarities in myeloid activation niches in perivascular and perifollicular areas of human skin, consistent with a role for the PVN in immune surveillance in healthy human skin.

**Figure 5:**
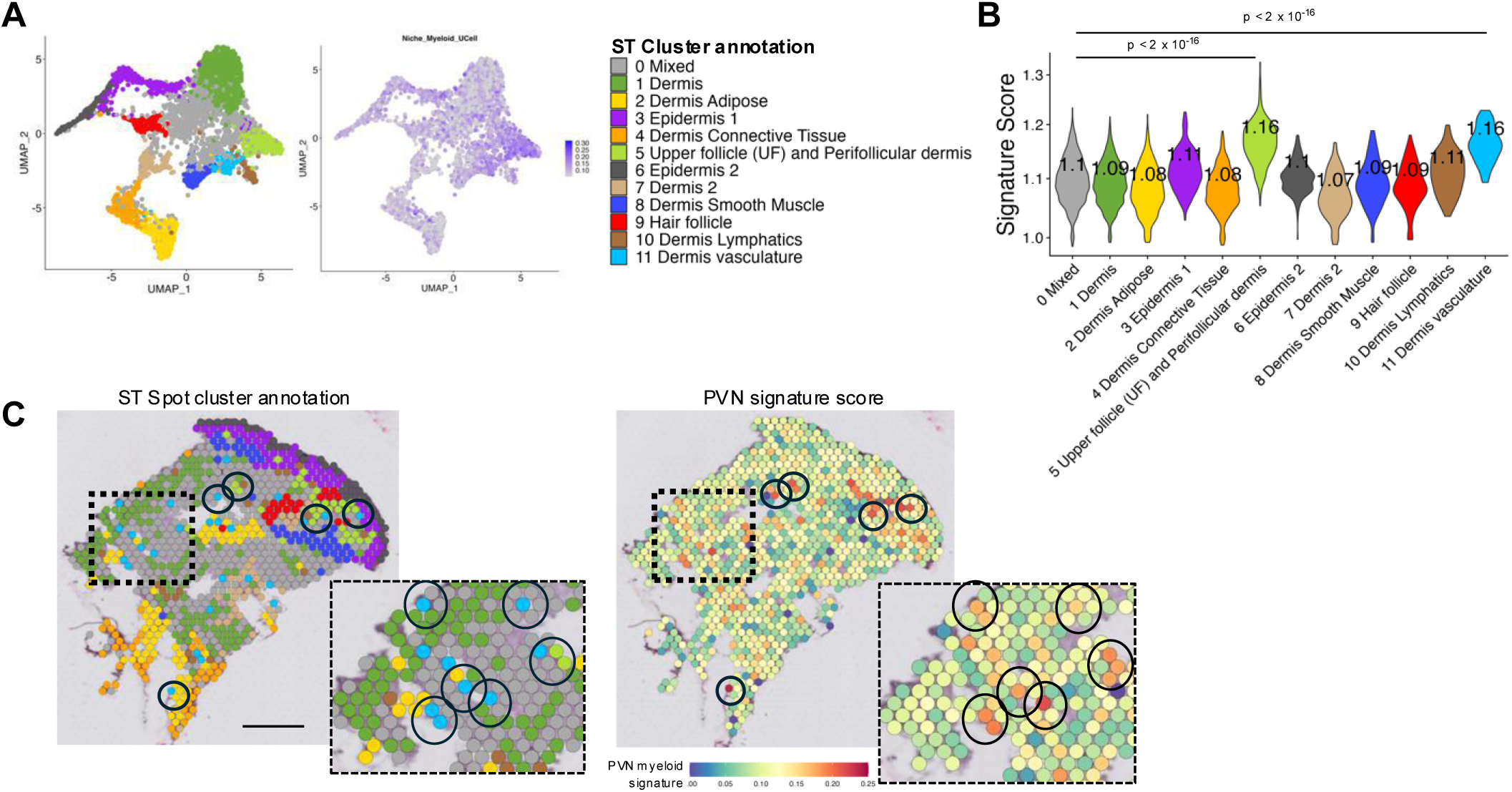
PVN-enriched myeloid activation signature identifies perivascular and perifollicular niches in healthy human skin. (A) UMAP visualization of PVN myeloid cell gene signature applied to spatial transcriptomic (ST) cell clusters from healthy human skin samples (*N* = 3, *n* = 5). (B) Quantitation of PVN myeloid gene signature by ST cluster. Statistical analysis performed using a pairwise t-test comparing cluster levels. (C) Representative tissue slice of healthy human skin, ST cluster spot distribution (left) according to previously annotated clusters and corresponding PVN gene signature score (right). Circles mark dermal vasculature spots (light blue) and/or perifollicular spots (light green) and corresponding enriched PVN signature scores, insert enlarged to show corresponding anatomical and PVN signature-enriched spots. Scale bar: 440 μm.

### CCR2-dependent APCs essential for Th1 activation in the PVN

Although the requirement of CD4 T-cell:APC encounter in peripheral tissues has been well-documented^41–47^, the specific APC subsets supporting such activation are poorly understood and findings are often difficult to interpret given many markers thought to define classic DCs, such as CD11c, are also upregulated on monocytes and macrophages within inflamed tissues. The main MHC-II^+^ subsets enriched in the niche were monocytes and cDCs (Figure 3E). To take a closer look at the APC potential of DCs and monocytes in the niche, we first applied an Ifng/Gbp/antigen presentation module transcriptional signature identified by comparison across various mouse infectious and inflammatory diseases^48^. Analysis revealed that many of the IFNγ/antigen presentation genes were enhanced in monocytes compared to DCs in the same location, and further elevated in the PVN (Figure 6A). To quantitate this finding, we applied the IFNγ/antigen presentation transcriptional signature to each cell type, revealing that niche-specific monocytes have an enriched IFNγ/antigen presentation signature compared to niche-specific DCs (Figure 6B).

**Figure 6:**
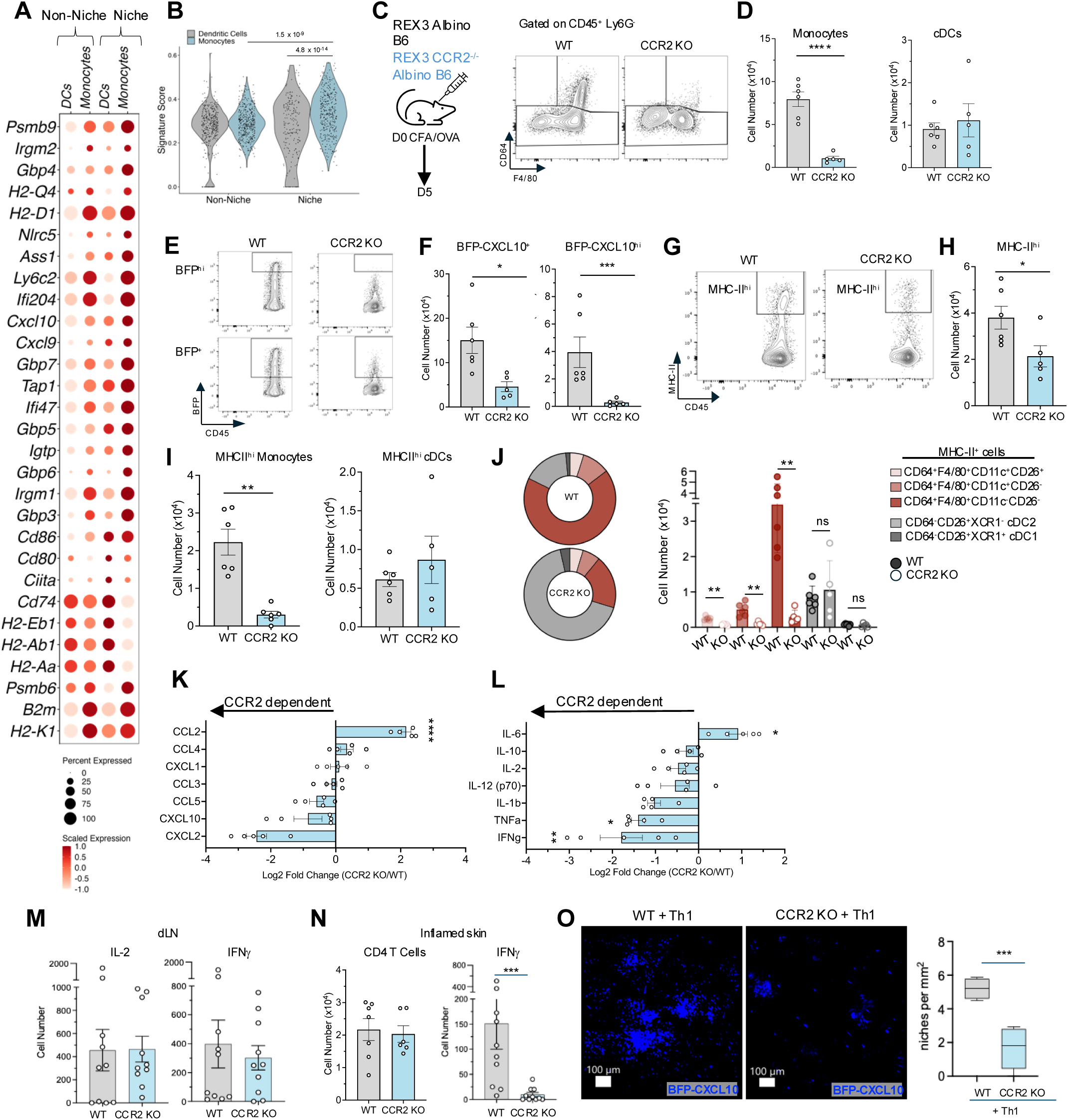
CCR2-dependent APCs are essential for Th1 activation in the skin. (A) Expression of IFNγ response-related genes within monocyte and dendritic cell clusters within Niche and Non-Niche regions. (B) IFNγ-response related gene signature applied to monocyte and dendritic cell clusters in niche and non-niche regions. Statistical analysis performed using a pairwise T-test comparing group levels. (C-J) REX3 WT and REX3 CCR2 KO mice immunized with OVA/CFA and flow cytometry analysis of the inflamed ear pinna performed on d5 p.i. (C) (left) Cartoon of experimental design for OVA/CFA immunization of REX3 WT and REX3 CCR2 KO mice, (right) CD64^+^F4/80^+^ monocytes in the inflamed dermis. (D) Number of monocytes (CD64^+^F4/80^+^) and cDC (CD64^−^F4/80^−^MHC-II^+^CD11c^+^CD26^+^) cells. (E) Representative plots of CD45^+^ cells gated on BFP-CXCL10^+^ and BFP-CXCL10^hi^. (F) Number of BFP-CXL10^+^ and BFP-CXCL10^hi^ cells. (G) Representative plots of MHC-II expression by CD45^+^ cells, gating on MHC-II^hi^ cells. (H) Number of total MHC-II^hi^ cells. (I) Number of MHC-II^hi^ monocytes and cDC cells. (J) Phenotypic analysis of monocyte and cDC populations in the inflamed skin of WT and CCR2 KO mice. (K, L) Luminex multiplexed bead-based immunoassay performed on d5 OVA/CFA ear lysates. Log2 fold change from the fold change comparing CCR2 KO to WT lysates. (M, N) WT and CCR2 KO mice immunized with OVA/CFA and skin-draining LN and ear cells isolated d5 p.i.. IFNγ and IL-2 ELISPOTS performed on cells from LN (M) and ear (N) in the presence of OVA peptide; (N) (left) CD4 T cell numbers determined via flow-cytometry, (right) IFNγ ELSPOTS/ear. (O) REX3 WT and REX3 CCR2 KO mice were immunized with OVA/CFA and 5×10^6^ OTII Th1 cells adoptively transferred 48-72h prior to analysis on d5 p.i.; (left) representative IV-MPM images and (right) number of niches per mm^2^. (D-J) Each dot represents an individual mouse and is shown as mean +/− SEM. Statistical analysis performed using Mann-Whitney test; *p<0.05, **p<0.01, ***p<0.001, ****p<0.0001. (D-J, M, N) Data from five independent experiments. (K, L, O) Data from two independent experiments.

To examine the role of monocytes in regulating Th1 activation in the PVN, we used CCR2-deficient mice^49^ crossed to the REX3 reporter. Following immunization with OVA/CFA, monocyte (CD64^+^F4/80^+^), but not cDC (CD64^−^CD11c^+^CD26^+^), recruitment to the inflamed dermis was severely compromised (Figure 6C, 6D). As predicted by our Niche-flow analysis (Figure 3E), blockade of monocyte recruitment led to a striking loss of CXCL10 expressing cells in the inflamed dermis, particularly the CXCL10^hi^ expressing cells (Figure 6E, 6F). The overall number of MHC-II^hi^ cells was also reduced (Figure 6G, 6H), attributed to a selective loss in MHC-II^hi^ monocytes but not a loss of MHC-II^hi^ cDCs (Figure 6I). Further phenotypic analysis revealed the MHC-II^+^ cells in the inflamed skin were predominantly CD64^+^F4/80^+^ monocytes, that were heterogeneous with respect to both CD11c and CD26 expression (Figure 6J, WT pie). This phenotyping suggests the monocyte population contains MHC-II^+^ moDCs (CD11c^+^CD26^−^), MHC-II^+^ moMacs (CD11c^−^CD26^−^) and a population of putative MHC-II^+^ inflammatory cDC2s (CD11c^+^CD26^+^)^50,51^. Notably, all these MHC-II^+^CD64^+^F4/80^+^ APCs were CCR2-dependent, being significantly reduced in CCR2-KO mice (Figure 6J), in contrast to cDC populations that were unchanged in the CCR2-KO (Figure 6J). Immuno-protein profiling of WT and CCR2 KO inflamed skin enabled the identification of CCR2-dependent chemokines and cytokines, with a specific loss of CCL5, CXCL10 and CXCL2 (Figure 6K). What was unexpected was a significant loss in Th1 effector cytokines IFNγ and TNFα in the absence of CCR2-dependent APCs (Figure 6L), even though the number of MHC-II^hi^ cDCs that could serve as APCs were similar in WT and CCR2 KO mice (Figure 6J).

To determine the role of CCR2-dependent APCs in supporting Th1 function, we examined the differentiation, recruitment and effector function of endogenously generated Th1 cells in WT and CCR2 KO mice following OVA/CFA immunization. While monocytes have been shown to traffic to inflamed LNs or spleen and contribute to T cell priming^52–55^, the absence of CCR2-dependent APCs did not alter the ability to prime and generate IL-2 and IFNγ producers in the dLN in response to OVA/CFA immunization (Figure 6M). Analysis of Th1 cells within the inflamed dermis, revealed that while CD4 T cell recruitment was not changed by the absence of CCR2-dependent APCs (and the corresponding loss of CXCL10 signals) (Figure 6N, left panel), the ability of those cells to secrete IFNγ was severely compromised (Figure 6N, right panel). These results highlight a critical requirement for CCR2-dependent APCs in peripheral Th1 activation in the inflamed skin. We had previously shown that the transfer of exogenously generated antigen-specific Th1 cells was able to enhance PVN numbers and size in an IFNγ-dependent manner^12^. Therefore, to further determine the biological impact of CCR2-dependent APCs on Th1 activation, we transferred OT-II Th1 cells into OVA/CFA immunized REX3 and REX3 CCR2 KO mice and used IV-MPM to assess Th1-driven PVN enhancement. For unbiased analysis of niche number, we employed a custom semi-automated density-based spatial clustering of application with noise (DBSCAN) algorithm, as previously described^12^. Consistent with the loss of Th1 peripheral activation in the absence of CCR2-dependent APCs, Th1 cells failed to enhance the number of PVN in the inflamed skin of CCR2 KO mice (Figure 6O).

### Intra-tissue assembly of the PVN driven by Th1 cells themselves

To determine the functional impact of Th1 activation in the PVN, we examined the Niche-seq T-cell cluster and conducted a differential analysis between the whole tissue, non-niche, and niche regions (Figure 7A). Niche-specific T-cells were enriched for genes associated with signal transduction and signal regulation in support of the PVN supplying early T cell activation signals. Interestingly, we did not see a consistent association with niche location and upregulation of effector cytokines, likely reflecting that Th1 cells are initially activated in the perivascular space within 24h of tissue entry but move into the tissue parenchyma to perform effector functions (Figure 2). Unique to the niche, was the marked enrichment of myeloid-recruiting chemokine gene expression by T cells in the PVN, compared with those T cells outside of the niche. These results suggested that activation within the niche may license T cells to assemble the niche itself.

**Figure 7:**
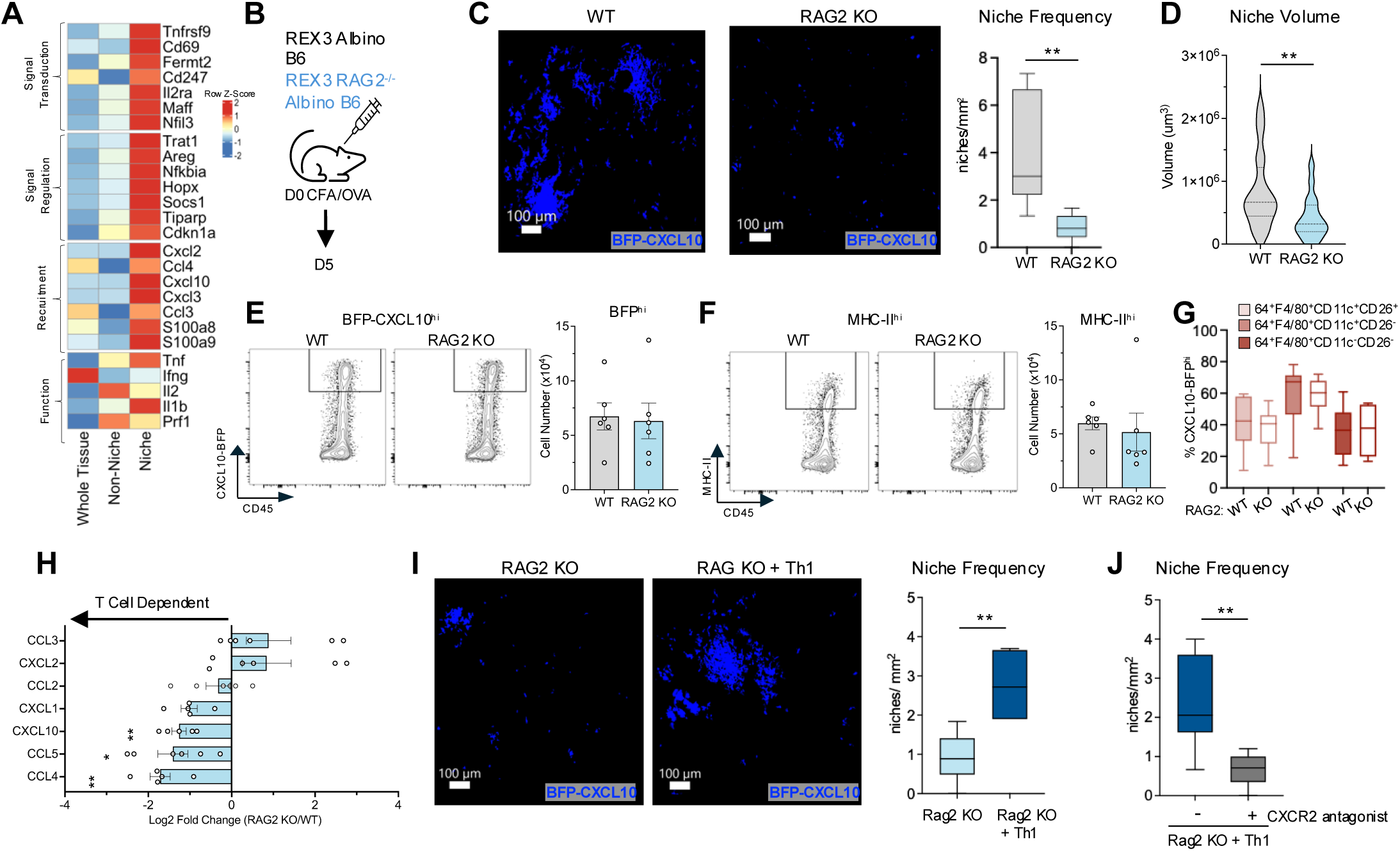
T cell licensing in the PVN enables Th1 cells to assemble the PVN by intra-tissue recruitment of myeloid cells. (A) Niche-enriched T cell signature derived from leading edge of GSEA. (B) Cartoon of experimental design for OVA/CFA immunization of REX3 WT and REX3 RAG2 KO mice with analysis of inflamed ear performed on d5 p.i.. (C) (left) Representative IV-MPM images in REX3 WT and REX3 RAG2 KO mice, (right) number of niches per mm^2^. (D) Niche 3D volume via DBSCAN and Imaris surface tracing. (E) (left) Representative plots of CD45^+^ BFP-CXCL10^hi^ cells, (right) number of BFP-CXCL10^hi^ cells. (F) (left) Representative plots of CD45^+^ MHC-II^hi^ cells, (right) number of MHC-II^hi^ cells. (G) Frequency of BFP-CXCL10^hi^ expressing cells among CCR2-dependent myeloid subsets. (H) Luminex multiplexed bead-based immunoassay of ear lysates from d5 OVA/CFA immunized mice. Log2 fold change was calculated comparing RAG2 KO to WT ears. (I) REX3 WT and REX3 RAG2 KO mice were immunized with OVA/CFA and 5×10^6^ OTII Th1 cells adoptively transfer 48-72h prior to analysis on d5 p.i., (left) representative IV-MPM images of BFP-CXCL10^+^ cells in the dermis, (right) number of niches per mm^2^. (J) Transfer of Th1 cells to REX3 RAG2 KO as in (I) with vehicle (-) or the CXCR2 inhibitor, SB225002 (+). (C, I) Each dot represents a single image covering a minimum of 1mm^2^ and is shown as mean +/− SEM. (E-H) Each dot represents an individual mouse and is shown as mean +/− SEM. (C-I) Statistical analysis performed using Mann-Whitney test; *p<0.05, **p<0.01, ***p<0.001. (D-I) Data shown is representative of three independent experiments. (H-J) Data shown are representative of two independent experiments.

To explore the contribution of T cells to the formation and/or maintenance of the CXCL10^+^ PVN, we first asked if T cells were required for the generation of PVN by using RAG2 KO mice crossed to the REX3 reporter. With the predicted cooperative communication network between myeloid cell subsets (Figure 4), we anticipated that myeloid cells would be sufficient to self-assemble into the CXCL10^+^ PVN on immune challenge. However, the loss of T cells led to a marked attenuation in the number and size of niches in the inflamed dermis (Figure 7B-D), consistent with earlier studies with acute CD4 T cell depletion^12^. To determine if this was a failure to recruit myeloid cells to the inflamed skin or an inability to upregulate CXCL10-BFP gene expression in the absence of T cells we assessed the immune infiltrate in the inflamed skin of REX3 WT and REX3 RAG2 KO mice. We found no defects in the recruitment of the major PVN-defining myeloid cell types to the inflamed skin of REX3 RAG2 KO mice.

The upregulation of CXCL10 and MHC-II expression associated with OVA/CFA was also unaffected by the absence of T cells (Figure 7E, 6F). Moreover, the dominant CXCL10^hi^ CCR2-dependent populations were also all present in the inflamed tissue of both WT and RAG2 KO mice (Figure 7G). Therefore, in the absence of T cells, myeloid recruitment to the inflamed skin and expression of CXCL10 was unaltered, but their ability to cluster into PVNs was severely compromised. Loss of clustering ability was associated with a reduction in the available myeloid-recruiting chemokines CXCL1, CXCL10, CCL5 and CCL4 in the inflamed skin in the absence of T cells (Figure 7H). The data suggested that T cells directly, or indirectly, impact chemotactic signals that organize myeloid cell intra-tissue aggregation within the inflamed skin. To test this idea, we asked if Th1 cells were sufficient to assemble tissue-recruited myeloid cells into niches. REX3 RAG2 KO mice were immunized with OVA/CFA to induce tissue recruitment of myeloid cells diffusely distributed across the dermal parenchyma. On d3 post-immunization, OT-II Th1 cells were acutely transferred and 48-72h later IV-MPM was used to assess niche assembly. Th1 cell add-back was sufficient for rapid intra-tissue assembly of CXCL10^+^ myeloid cells into perivascular niches (Figure 7I). T cells in the niche were enriched for several chemokines that can drive monocyte and DC migration, including the CXCR2 ligands Cxcl2 and Cxcl3. Indeed, the ability of Th1 cells to assemble the PVN was significantly impaired following administration of the CXCR2 inhibitor, SB225002 (Figure 7J)^56^.

In the early steps of region immunity in inflamed tissues innate immune cell clustering has been demonstrated to be orchestrated by cooperative signaling between myeloid cell subsets, namely the cross-talk between neutrophils and monocytes or macrophages^16,57^. Here we reveal a new positive feedback loop at the time adaptive immune cells join the inflamed tissue, where the incoming effector T cells themselves play a key role in the intra-tissue repositioning of myeloid cells to cluster at perivascular sites. These perivascular niches provide the first opportunity for effector Th1 cells to encounter antigen as they enter the inflamed skin and also provide instructional signals that impart Th1s with tissue organizer functions that catalyze niche assembly.

## DISCUSSION

The presence of perivascular infiltrates in non-lymphoid peripheral tissues has been observed across numerous inflamed, infected and malignant tissues^2^. Using high-resolution dynamic imaging or spatial transcriptomic techniques these sites have emerged as critical immune niches for modulating T cell effector function. Here we integrated dynamic time-resolved intravital multiphoton microscopy (IV-MPM) with spatially-resolved immune profiling to address fundamental questions about the spatiotemporal dynamics of Th1 cell activation following entry into the inflamed skin. We show that in the first 24h following tissue entry, Th1 cells are enriched in CXCL10^+^ perivascular niches (PVN) and express markers consistent with niche-restricted antigen-specific activation, providing new insight into the location and timing of initial activation of Th1 cells in the inflamed skin. PVN residency was transient, with the majority of Th1 cells moving from the PVN into the tissue parenchyma within 24h of tissue entry. Spatially-resolved single-cell interrogation of the PVN revealed a site-specific myeloid hub with a transcriptional signature that highlights cooperative activation programs between myeloid cell subsets and locally enriched chemokine production. The scRNAseq data enabled the development of a myeloid PVN transcriptional signature that was sufficient to identify similar perivascular and perifollicular regions of myeloid cell activation in healthy human skin. Mechanistically, the niche-enriched CXCL10^+^ myeloid cells present in the inflamed skin originated primarily from CCR2-dependent recruitment of inflammatory monocytes, and those inflammatory monocytes were critical for Th1 activation in the skin. In turn, Th1 cells received signals within the PVN that licensed them for niche assembly, facilitating the intra-tissue repositioning of the CXCL10^+^ myeloid cells to the PVN in a CXCR2 dependent manner. Our work highlights the conceptual advances that can be made by integrating dynamic, temporal and spatial single cell analytical tools, uncovering a complex positive feedback loop within the perivascular space that regulates myeloid and T cell positioning for the exchange of critical signals that optimize early T cell activation upon tissue entry. Notably, it was the incoming Th1 cells themselves that were responsible for the nucleating myeloid cells into perivascular immune hubs, adding a tissue organizer role to their ‘helper’ functions in peripheral tissues.

The clustering of myeloid cells within microanatomical niches is thought to enhance their capacity to mount effective immune responses by facilitating critical cell-cell interactions that upregulate production of pro-inflammatory cytokines and chemokines and activate anti-microbial functions and antigen-presenting capacity. Our analysis identified not only the spatially-restricted upregulation of chemokine-mediated recruitment pathways within these niches but also common myeloid cell activation pathways, suggesting that myeloid cells acquire distinct functional properties in the PVN, compared to their parenchyma counterparts. This was particularly apparent when comparing IFNγ− and IFNβ-responsive signature scores across myeloid subsets, with all niche resident populations enriched for IFNγ-responsive programming. Interestingly, *in vitro* studies suggest chemokine-dependent clustering of monocytes facilitates monocyte differentiation into monocyte-derived DCs and macrophages^58^, thus the predominance of monocyte-derived APC in the PVN maybe due to in situ differentiation of newly recruited monocytes within the niche itself. Consistent with our own identification of shared activation programs across myeloid cell subsets in the PVN, spatial transcriptomics studies of human glioblastoma and glioma have demonstrated that myeloid cells with similar programs preferentially cluster together, driven by strong migratory cues from other immune cells^59^.

Our niche-seq analysis revealed that the monocyte cluster within the PVN exhibited a heightened responsiveness to IFNγ compared to the dendritic cell (DC) cluster, and an upregulation of MHC and costimulatory machinery for APC function. Indeed, blocking monocyte recruitment to the inflamed dermis led to a significant reduction in IFNγ levels, and a striking absence of Th1 effector function. Combined, these results indicate a critical positive feedback loop between monocytes and T cells that drives dermal inflammation. Our studies support the notion that CCR2-dependent APCs (monocytes and/or inflammatory cDC2s) recruited to peripheral tissues are the primary APCs for Th1 activation in the dermis, and show that their positioning in the PVN correlates with heightened activation and responsiveness to IFNγ. This local, microanatomical, cross-talk between monocytes and Th1 cells may be critical for honing monocyte APC function. Many studies have implicated a role for moDCs in supporting Th1 and CD8 T cell activation in peripheral tissues^60–64^. More recently, the identification of inflammatory cDC2s that are often phenotypically similar to moDCs (expressing CD11b, Ly6C and CD64, and partially dependent on CCR2)^50,51,65,66^, has complicated the interpretation of earlier studies implicating monocytes in peripheral T cell activation. Monocyte transfers in models of sterile injury, infection and autoimmune models^52,53,62,67,68^ provide a direct demonstration of monocytes (independent of inflammatory cDC2s) as pivotal APCs in peripheral tissues. How the PVN microenvironment ‘conditions’ CCR2-dependent APCs for Th1 activation will require further investigation. Similarly, there remains an important knowledge gap in our understanding of the specific signals that CCR2-dependent APCs provide to effector CD4^+^ T cells for peripheral activation, signals that appear insufficiently supplied by co-localized cDC populations in the same inflamed tissue. Monocyte:T cell clustering could directly activate Th1 cells, or their co-localization could supply local cooperative signals that support differentiation or feedback to enhanced cDC APC function^55^. Additionally, signals in the PVN may support effector T cell survival. In a cutaneous tumor model, cDC3’s expressing CXCL16 were implicated in promoting CD8 effector survival in the PVN^69^. We also find enhanced CXCL16 expression in our PVNs, enriched in both DC and monocyte clusters, with a corresponding upregulation of CXCR6 on incoming Th1 cells using our time-resolved model. Thus, the PVN may provide both initial activation and survival signals to Th1s upon skin entry.

In the past decade, the ability to visualize innate immune cell clustering in situ, in real time, with IV-MPM, has enabled dissection of the sequential recruitment and cooperative positioning of innate effectors early in response to tissue damage or infection (hours to days after the challenge)^70–72^. Dramatic neutrophil swarming behavior driven by an LTB4-dependent signal relay, precedes a neutrophil-dependent monocyte recruitment to the microanatomical clusters around sites of sterile injury^57^. In infection models, early immune clustering in infected tissues involves neutrophils and/or macrophages, with examples of each one of these subsets being the initial driver of immune aggregation^16,57,73,74^. Early NK-derived IFNγ has been implicated in orchestrating myeloid cell recruitment, positioning and activation for initial pathogen control^75,76^. These initial immune aggregates can rapidly disperse upon repair of tissue damage or control of the initial pathogen threat. In contrast, how immune aggregates are re-enforced or propagated later in the immune response as effector T cells enter peripheral tissues (3-6 days after the challenge) is less well understood. Our observations indicate that the Th1 cells themselves help to aggregate CXCL10^+^ myeloid cells into these perivascular activation niches. In the absence of T cells, skin-recruited myeloid cells were diffusely distributed in the inflamed dermis 5 days post-immunization. Reintroducing Th1 cells into the inflamed dermis was sufficient to restore the intra-tissue clustering of CXCL10^+^ myeloid cells at the PVN. The ability of Th1 cells to mediate niche assembly appears to be acquired upon tissue entry by initial activation in the perivascular space, through the spatially-restricted upregulation of CXCR2 myeloid-attracting chemokines. Thus, Th1 cells are crucial architects of the PVN, assembling a myeloid hub that supports their initial activation upon tissue entry. We have previously shown that IFNγ was important in PVN formation^12^, therefore we propose a temporal hand off of IFNγ sources from early NK to late Th1 cells in the shaping of myeloid cell aggregates that nucleate exchange of activation signals in inflamed tissues. While IFNγ can have direct effects on enhanced APC function, as we see with the PVN enriched IFNγ-responsive signatures in monocytes and DCs, data from a model of contact hypersensitivity (CHS) suggests local non-hematopoietic cells can also contribute to niche organization. In an oxazolone-induced CHS model, myeloid cell clustering occurs around hair follicles, and local T cell-derived IFNγ triggered keratinocytes to produce CCL2 that boosted monocyte clustering^77^.

The fundamental understanding of the role of PVNs in health and disease has been limited by a lack of consistency in the definition of these microanatomical regions and the inability to properly define the boundaries between perivascular and parenchymal compartments in different tissues^5^. Our studies have defined a myeloid PVN transcriptional signature that can identify regions of similar myeloid cell activation in healthy human skin that were also spatially restricted to perivascular and perifollicular regions^40^. Several cancer studies identify beneficial myeloid cell niches that help promote anti-tumor immunity and response to checkpoint therapy. Myeloid APC niches in ovarian cancer that facilitate CD28 costimulation can enhance the effector function of exhausted CD8+ T cells^78^, while DC-CD4 T cell niches in hepatocellular carcinoma boost CD8 function following PD-1 blockade^79^. A CXCL10^+^ ‘immunity hub’, enriched in vascular cells, in human lung cancer harbors stem-like T cells and is strongly associated with beneficial outcomes to PD-1 blockade^17^. Similarly, functionally distinct myeloid clusters can be predictive of patient survival in glioblastomas^59^. Moreover, in acute Graft-Versus-Host Disease, vascular myeloid niches have been associated with cases where steroid therapy was effective, highlighting their potential as prognostic markers^80^. Thus, our ability to identify transcriptionally distinct activation niches within tissues should lead to rapid advances in our understanding of the functional implications of perivascular immune clusters in the steady state and in disease states.

Compartmentalization of immune responses in space and time is fundamental to efficient tissue homeostasis and prompt immune activation upon microbial, environmental or mechanical challenge. Spatially constrained immune activation also limits collateral damage to healthy regions within the target tissue. Immune regulatory niches have been identified at points of pathogen entry at barrier surfaces, such as the hair follicles in the skin^81^ and the lamina propria of the gut^82^, and within tissues at points of interstitial drainage^7^. Upon pathogen challenge, these homeostatic niches can be transformed into sites of immune activation^82^, or new immune activation niches can be formed perivascularly at points of leukocyte entry into tissues^11,12^. We provide new temporal and spatial insight into the initial events in effector Th1 cell activation in the inflamed skin. In coming Th1 cells spend the first ∼24h in the skin in a CXCL10^+^ perivascular niche, coincident with initial antigen-specific tissue activation, before moving into the tissue parenchyma. Monocyte-dependent activation in the PVN licenses Th1s for tissue organizing function. Th1 cells were necessary and sufficient to amplify niche assembly through intra-tissue repositioning of CXCL10^+^ myeloid cells to the PVN. This new understanding of the PVN can inform strategies to enhance or attenuate regional immunity through target interventions and modified cell therapies.

## Supporting information

Bala, McGurk et al Supplementary Figures

## ACKNOWLEDGEMENTS

We thank the members of the Fowell lab for helpful discussions on the studies. Brian Rudd and Mandy McGeachy provided insightful comments on the manuscript. We gratefully acknowledge support from the following agencies: National Institutes of Health Grants P01 AI102851 and R37 AI072690 to DJF; T32 EB023860 to NB. We acknowledge the support of the Imaging (RRID:SCR_021741) and Flow Cytometry core (RRID:SCR_021740) facilities at Cornell University

## AUTHOR CONTRIBUTIONS

N.B., A.M., S.A.L., and D.J.F. designed the experiments. N.B., A.M., S.A.L., and E.M.C. performed the experiments. N.B. performed niche bioinformatics and computational analysis. I.S. and S.N. generated spatial transcriptomics data from healthy human skin and aided in the application of the PVN myeloid signature to the ST human data. N.B., A.M, S.A.L., and D.J.F. wrote the manuscript.

## SUPPLEMENTARY FIGURES

**Figure S1. Multiphoton microscopy-guided identification of niche regions for biopsy.** REX3 mice were immunized with OVA/CFA and IV-MPM preformed d5 post-immunization, IV-labeling of CD31 with Alexa 647 (pink) 30 mins prior to imaging. A) Spatial coordinates of reference points made on the outer edge of the ear pinna with permanent marker, as well as CXCL10-BFP PVN rich regions (niche-enriched) and CXCL10-BFP devoid regions (non-niche) by MPM were mapped with the Olympus user interface “register area” function. Using coordinates from each reference point and individual registered area, x (μm) and y (μm) distances were calculated and checked for agreement between reference points 1 and 2. Regions identified by these calculated distances were then biopsied (2 mm punch biopsy) and subsequently reimaged. B) Representative image of niche-enriched and non-niche 2 mm punch-biopsies. C) Flow cytometry of MPM guided biopsy material from niche-enriched and non-niche regions of the inflamed ear pinna, compared to random punch biopsies of the inflamed ear pinna.

**Figure S2. Spatially-resolved scRNAseq cell clustering and analysis.**

REX3 mice were immunized with OVA/CFA and IV-MPM preformed d5 post-immunization, with IV-labeling of CD31 with Alexa 647 (pink) 30 mins prior to imaging. Niche and non-niche biopsies, as well as whole tissue (ear pinna) were processed for scRNAseq. (A) Heatmap of top 10 genes represented in each cell cluster from UMAP representation. FindMarkers function in Seurat was used to determine marker genes for distinguishing each cluster from the remaining cluster. Row Z-score represents normalized expression across each gene. (B) Dotplot of two identifier genes per cluster used for annotation of each cluster identified within the UMAP. (C) Quantified cluster distribution percentages within the whole tissue. (D) Niche scored by responsiveness to type I IFN (IFN-β) or IFNγ and overlaid on UMAP. (E) IFNγ signature score in CXCL10^hi^ and CXCL10^lo^ monocytes in niche and non-niche regions.

**Figure S3. Sustain neutrophil recruitment is not required for CXCL10 induction or niche formation.**

(A) Quantification of neutrophil infiltrate in d2 and d5 CFA/OVA immunized ears following treatment with 200mg non-specific IgG isotype control mAb or anti-Ly6G mAb (1A8) and 400 mg anti-mouse kappa light chain mAb (MAR 18.5). Neutrophils were pre-gated on Live/CD45^+^/CD4^−^/CD64^−^/F4/80^−^/MHCII^−^ cells and identified as CD11b^+^ Ly6C^+^ cells, excluding Ly6C^hi^ monocytes. (B) Frequency of BFP-CXCL10^+^ cells in d5 CFA/OVA immunized ears, pre-gated on Live/CD45^+^ cells. (C) Quantification of niche formation in d5 CFA/OVA immunized ears per mm^2^. (A-C) Statistical analysis performed using Mann-Whitney test; *p<0.05

**Figure S4. PVN myeloid gene signature.** The PVN gene signature was compiled with genes enriched in the niche in 3 out of 4 myeloid cell populations: neutrophils, monocytes, macrophages and dendritic cells. In addition, niche-enriched genes associated with IFNγ and antigen presentation that were elevated in PVN monocytes.

## SUPPLEMENTAL VIDEO DESCRIPTIONS

**Video S1. Antigen-specific Th1 arrest in the perivascular niche.**

*In vitro* generated LCMV GP_61-80_-specific (SMARTA) Th1 OFP cells were intravenously co-transferred alongside ovalbumin-specific (OT-II) Th1 GFP cells into REX3 mice intradermally immunized with OVA/CFA in the ear pinna. Day 5 post-immunization IV-MPM was used to visualize the inflamed dermis and identify the CXCL10-BFP^+^ PVN. Distribution of BFP-CXCL10^+^ cells (blue) in the dermis and motility tracks of OT-II GFP^+^ (green) and SMARTA OFP^+^ (red). The video represents a 2D maximal z-projection time series of a 60μm-thick imaging volume; scale bar 80µm.

## STAR★Methods

### Key resources table

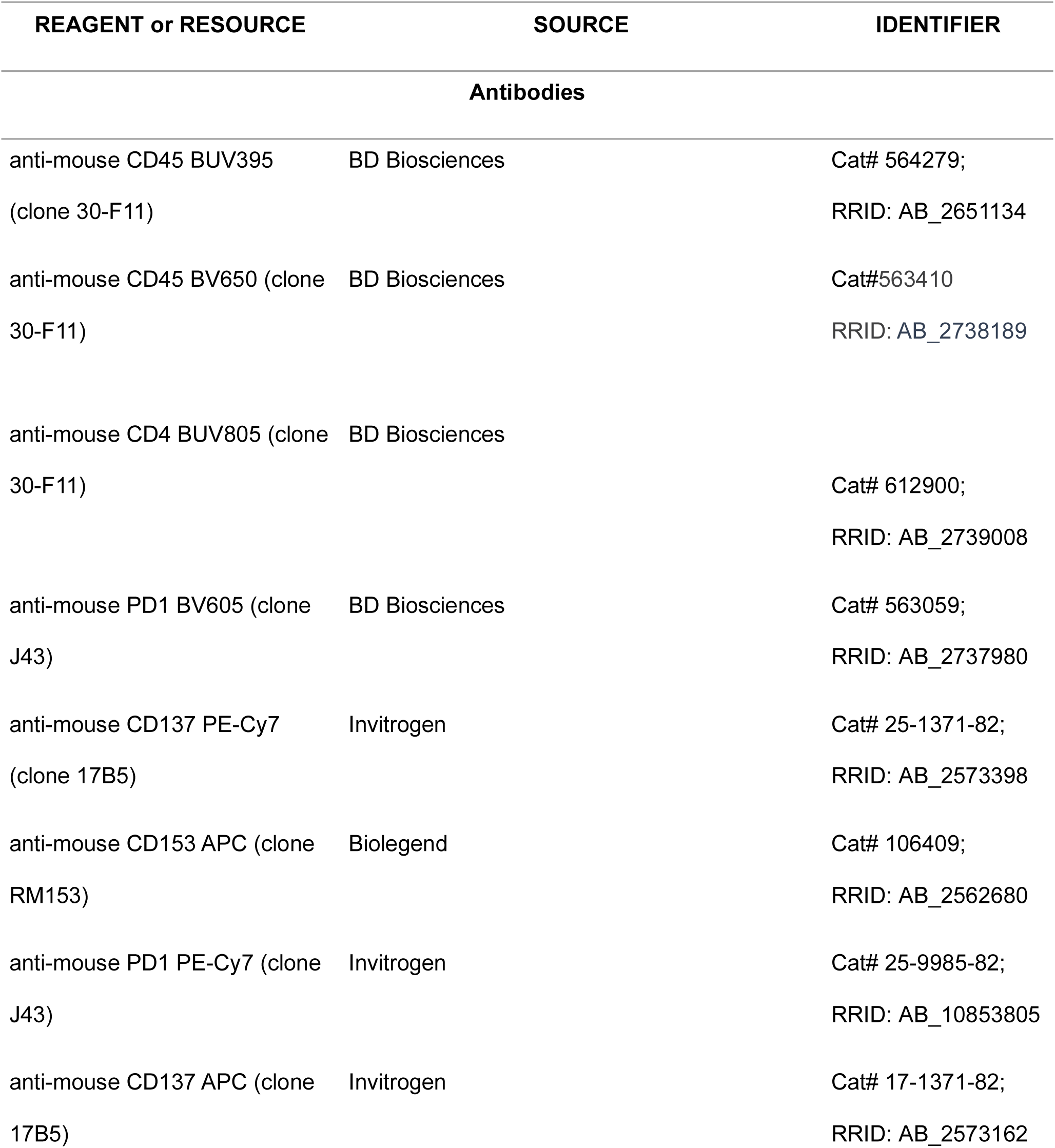
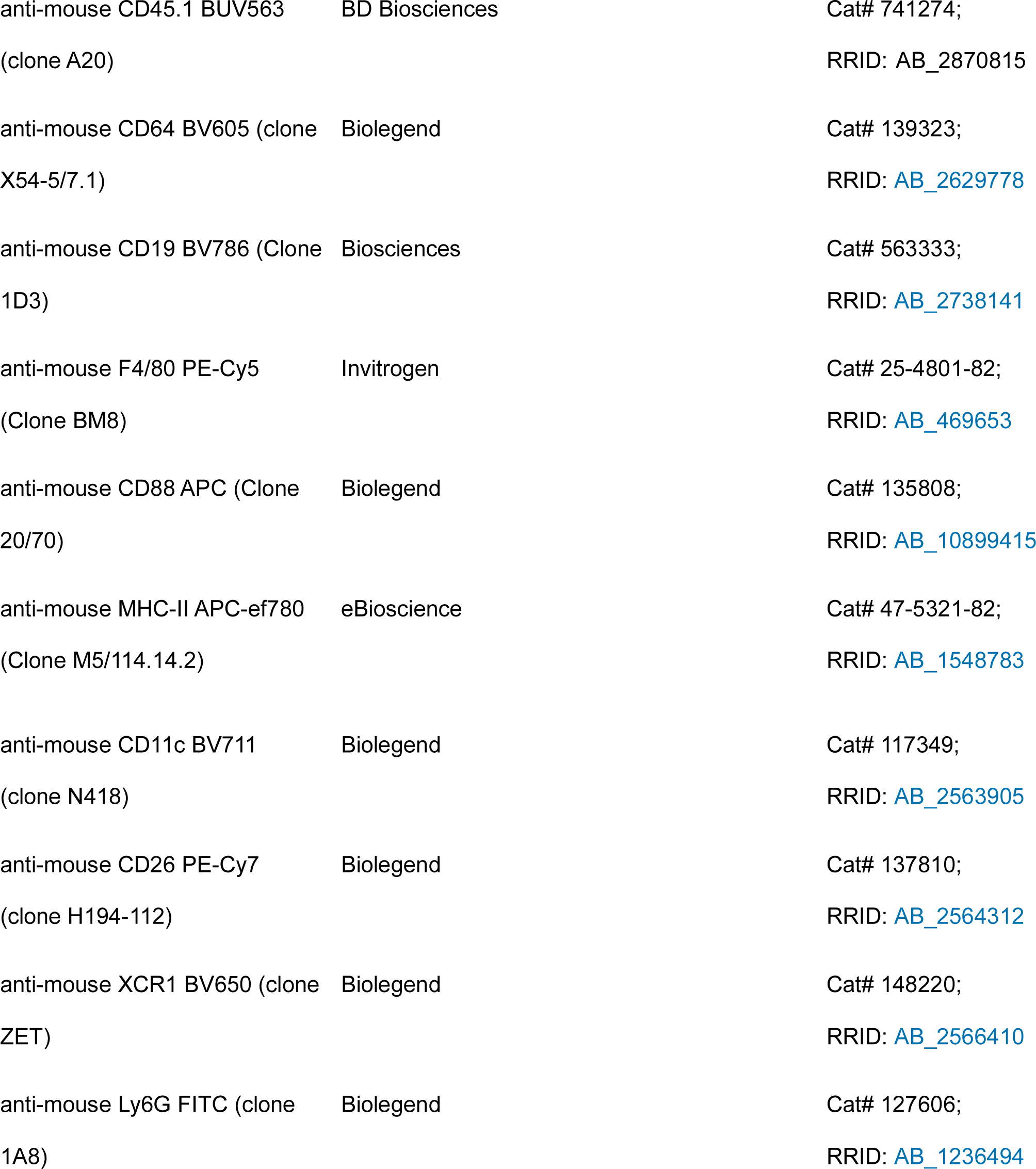
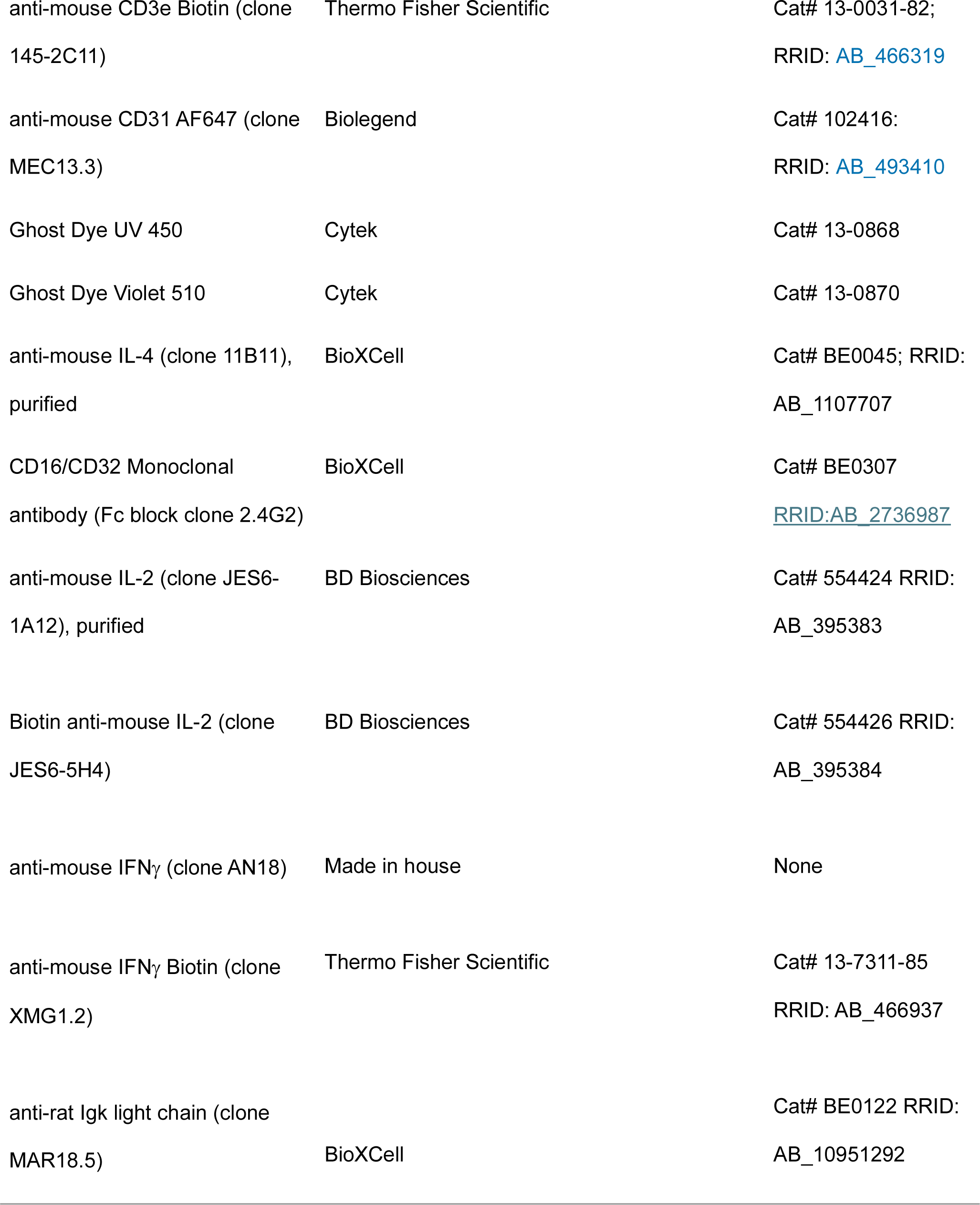
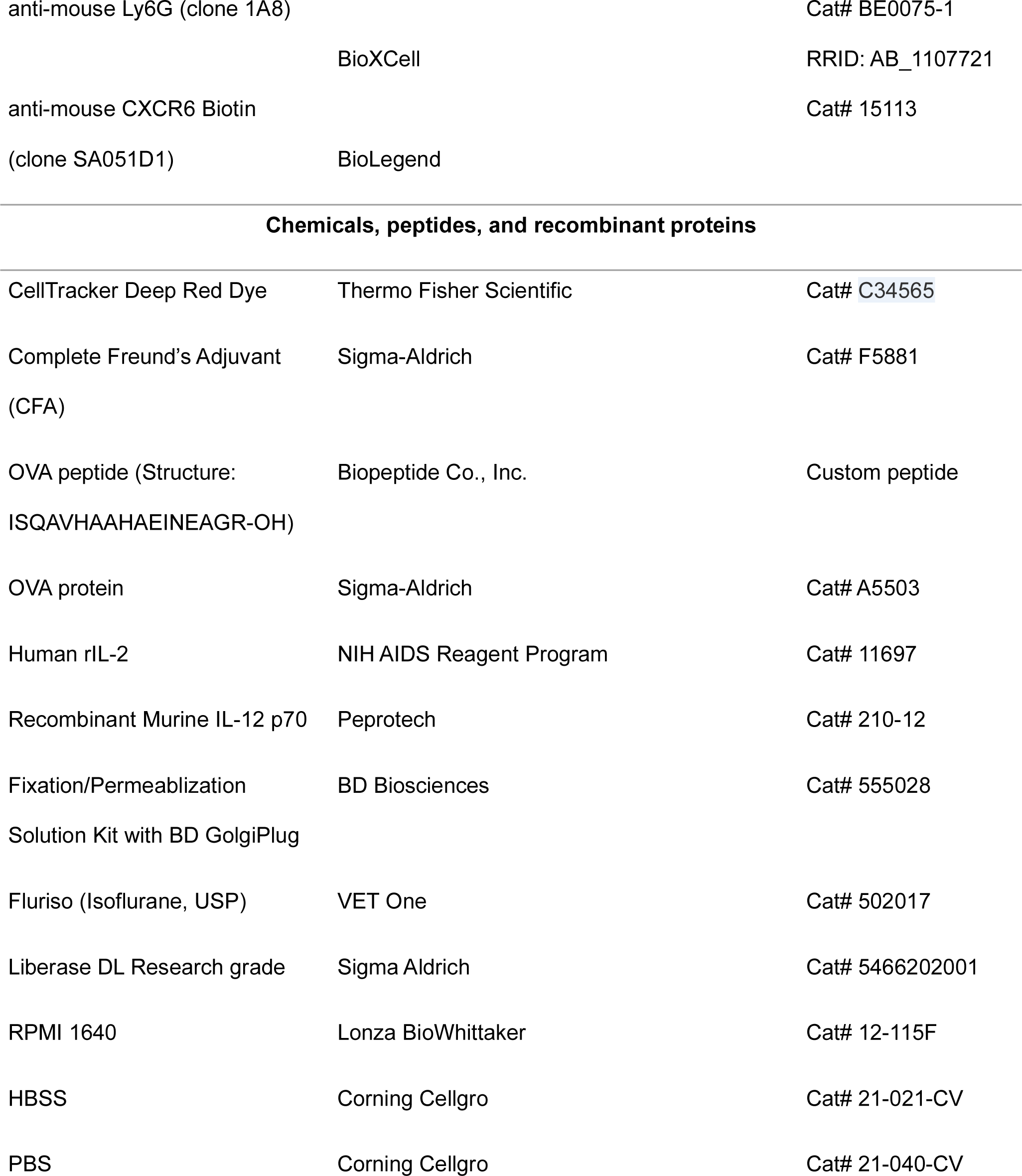
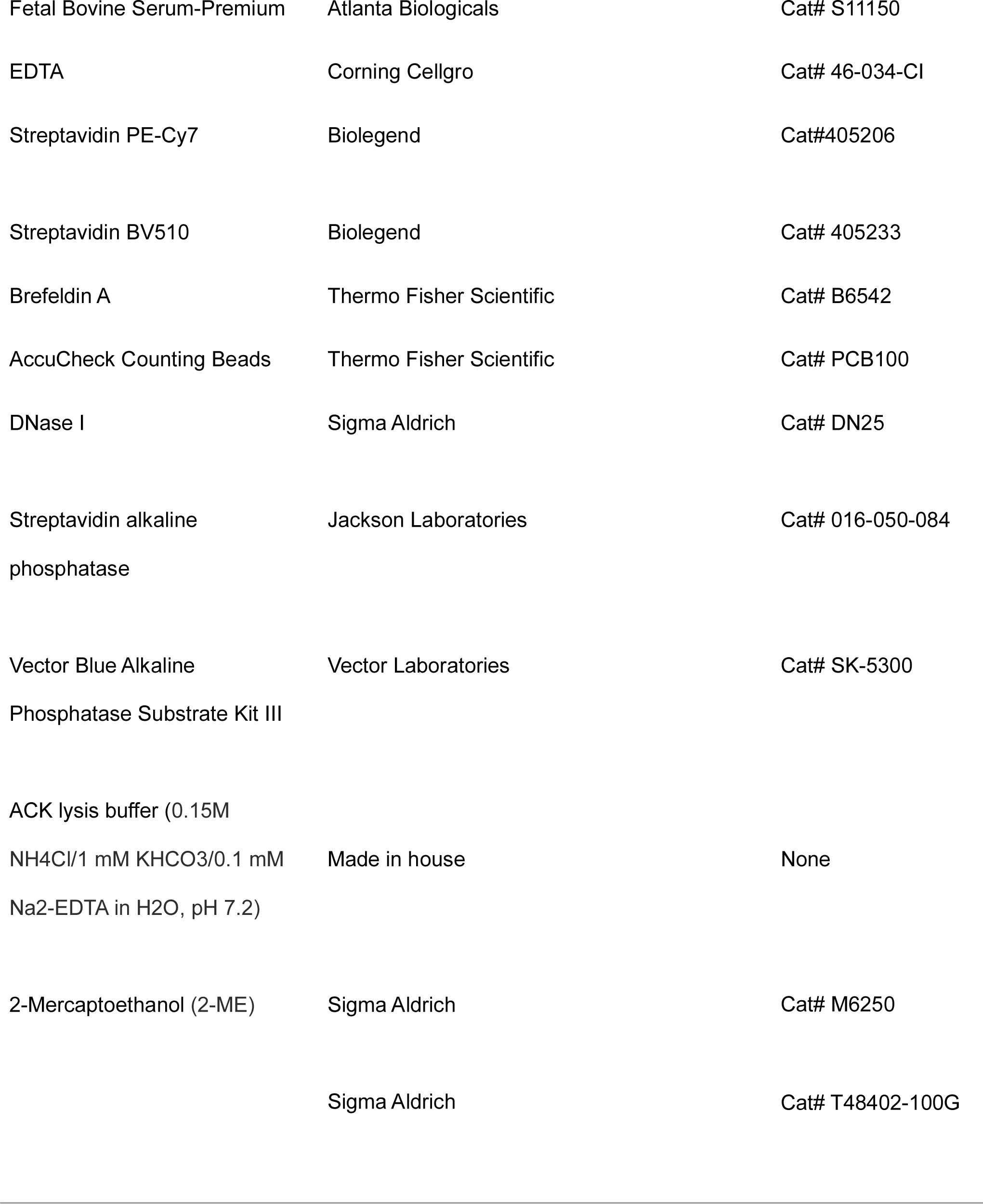
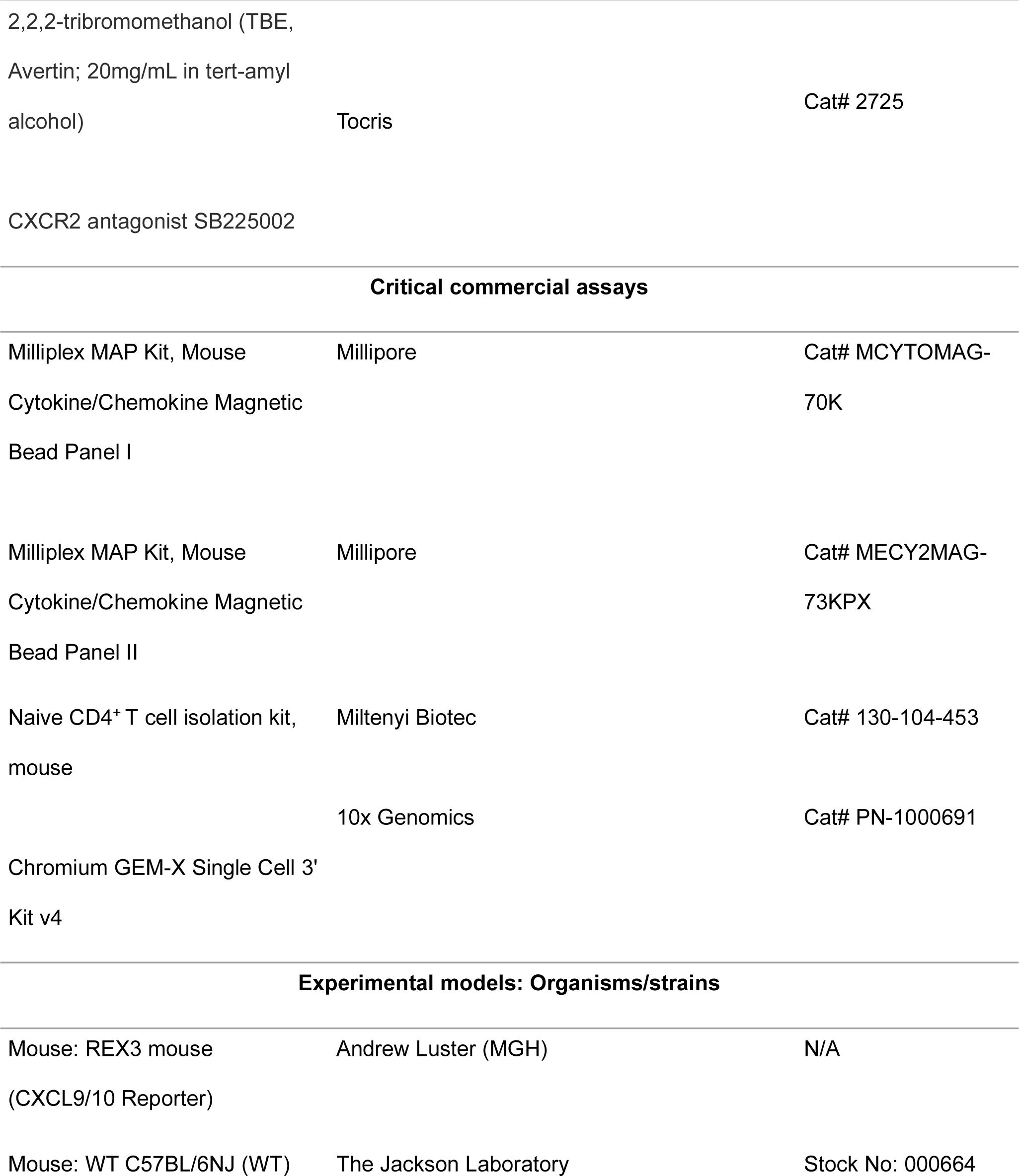
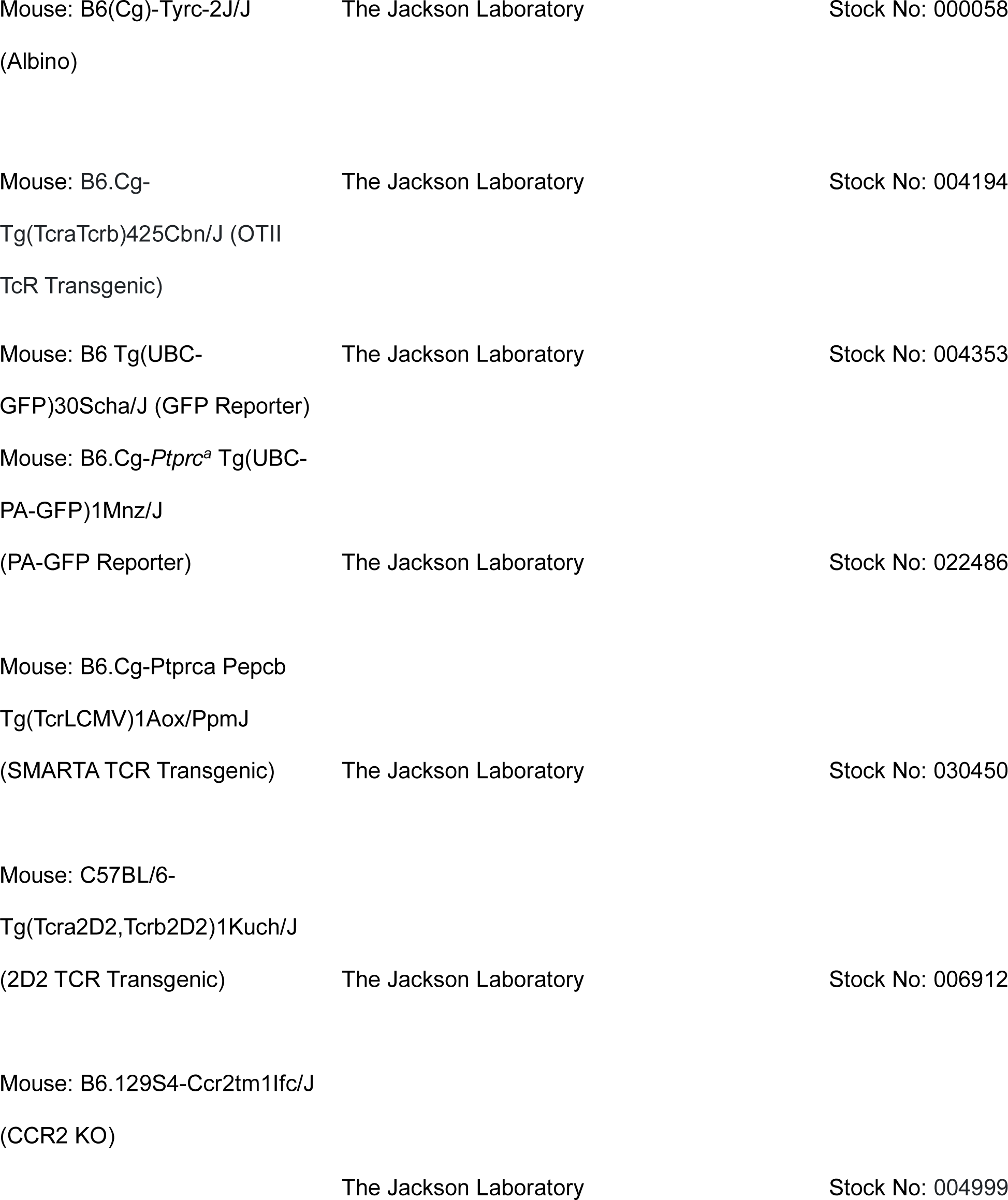
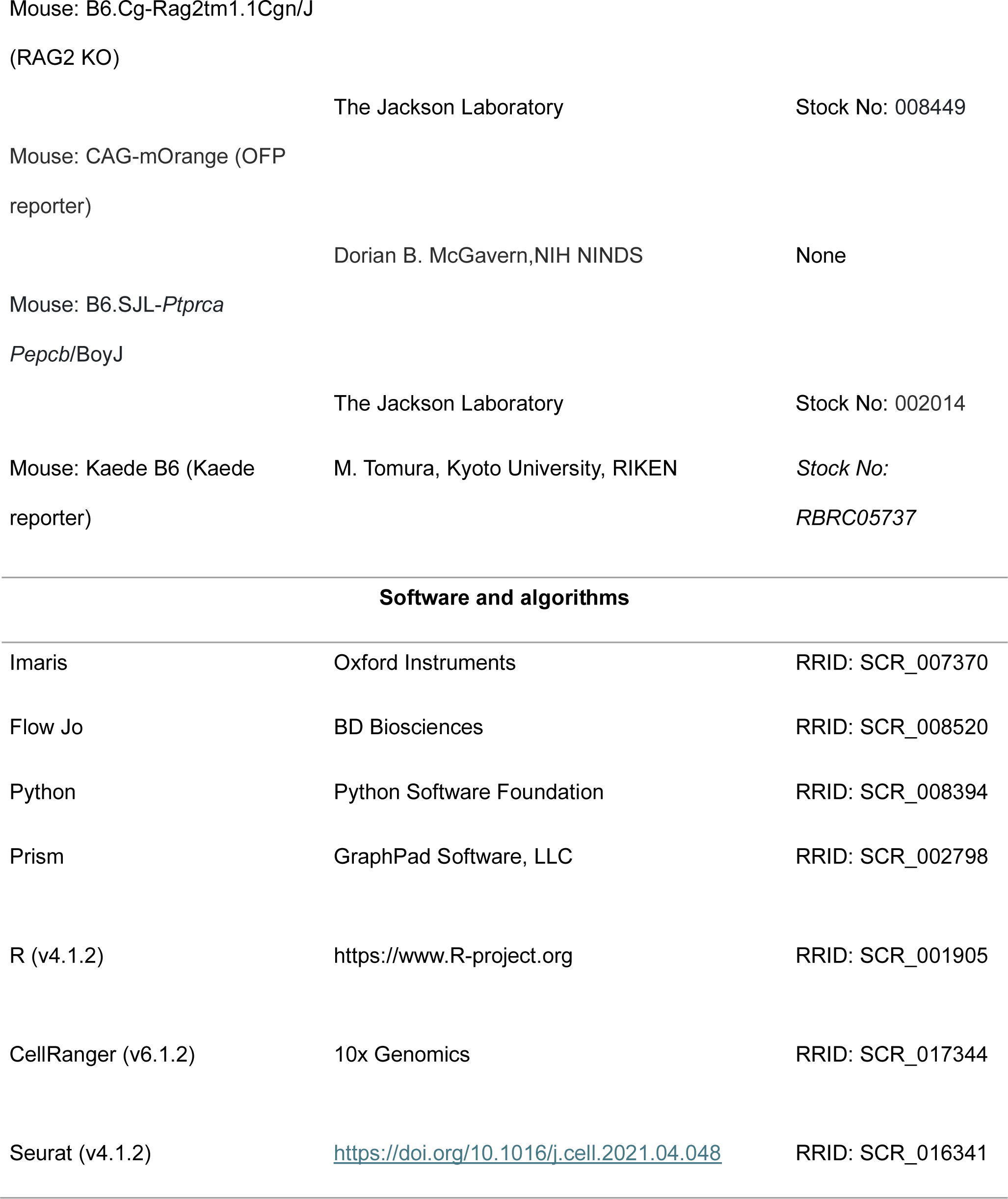
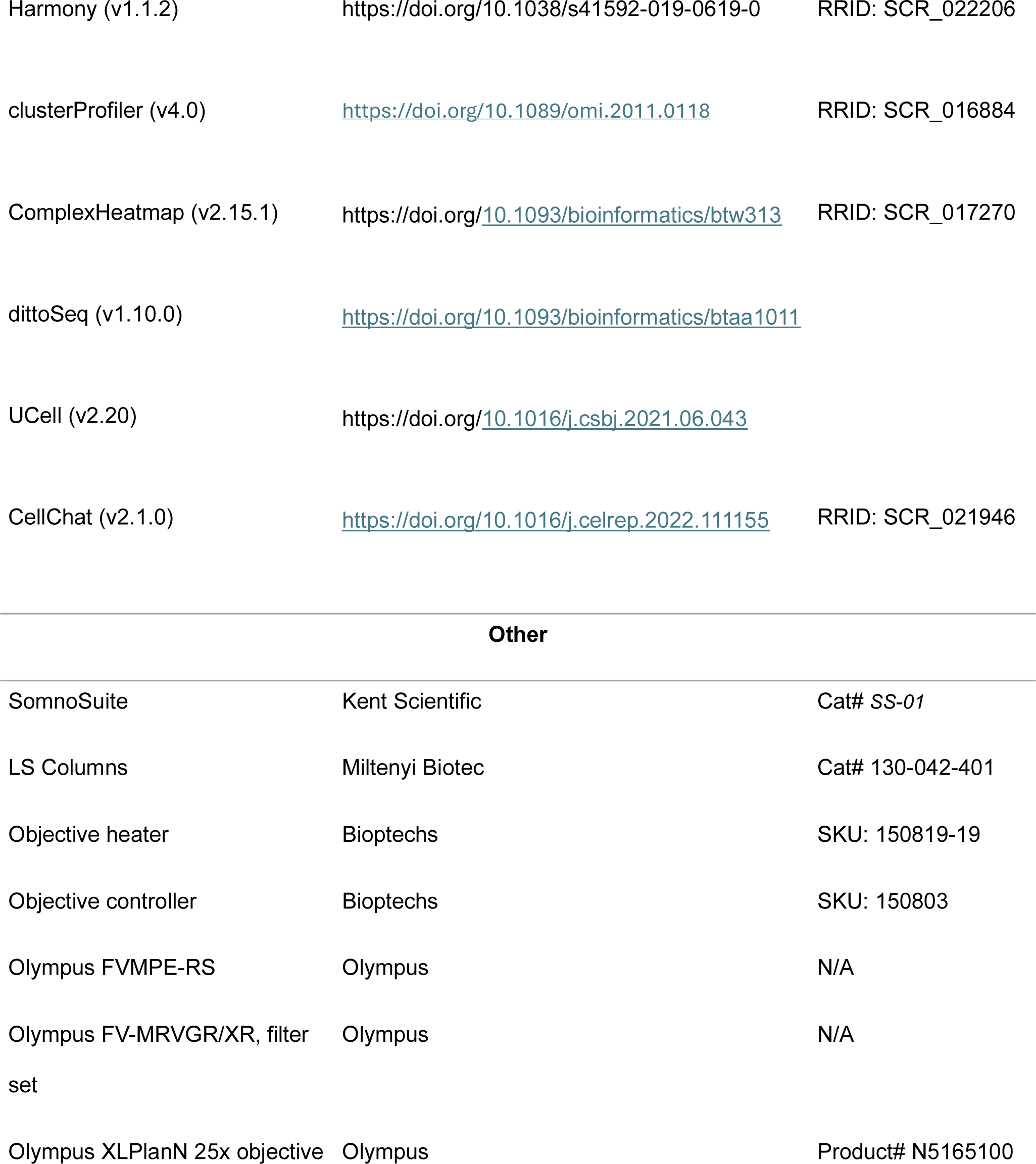
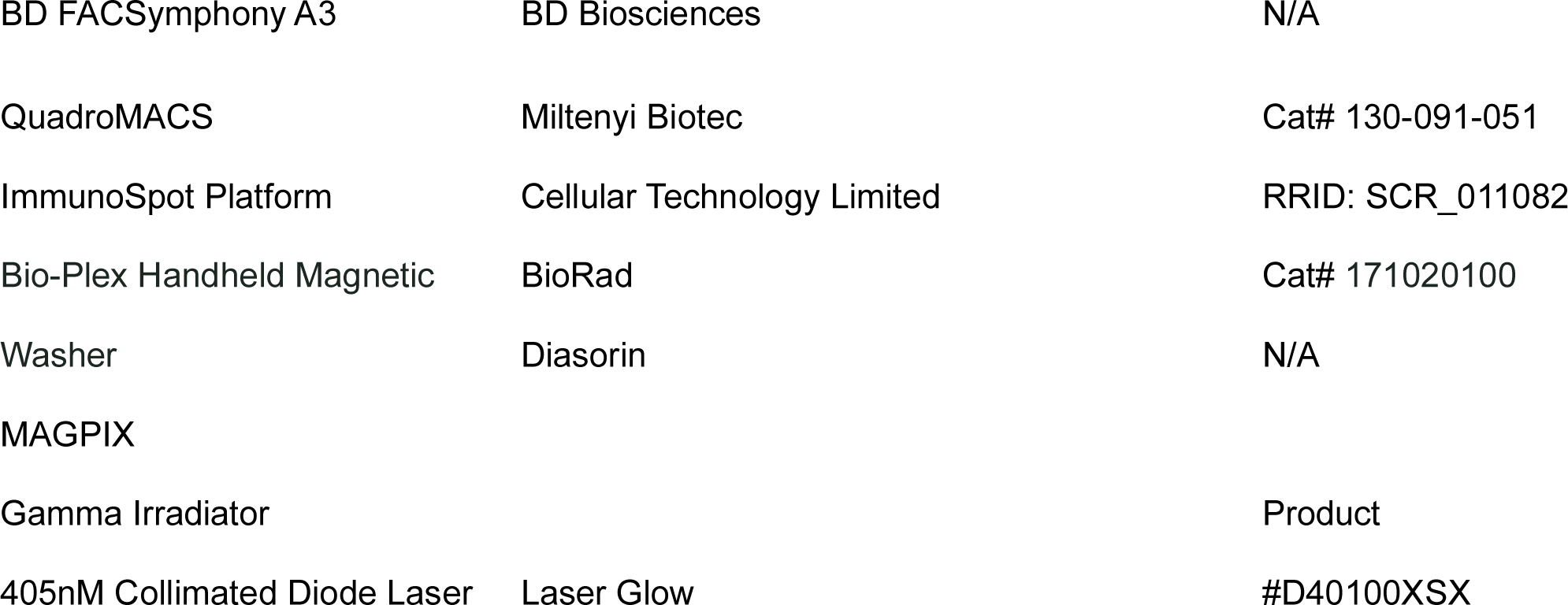

### Data and code availability

Raw data for scRNAseq will be deposited in the public database, GEO pending. Original code is available from the lead contact upon request.

### Lead Contact and Materials Availability

Further information and requests for resources and reagents should be directed to and will be fulfilled by the Lead Contact, Deborah J. Fowell.

### Experimental Model and Subject Details

All animal procedures were approved by the Institutional Animal Care and Use Committee of Cornell University. Both female and male mice were bred and maintained under pathogen-free conditions in group housing. Mice used in experiments were between 6 and 14 weeks of age and were euthanized in accordance with Cornell University guidelines. The following mouse strains were purchased from The Jackson Laboratory: C57BL/6NJ (WT), B6(Cg)-Tyrc-2J/J (Albino), B6.Cg-Tg(TcraTcrb)425Cbn/J (OTII TcR Transgenic), B6.SJL-*Ptprca Pepcb*/BoyJ (CD45.1 congenic), C57BL/6-Tg(UBC-GFP)30Scha/J (GFP Reporter), B6. B6.Cg-*Ptprc^a^* Tg(UBC-PA-GFP)1Mnz/J (PA-GFP Reporter), B6.Cg-Ptprca Pepcb Tg(TcrLCMV)1Aox/PpmJ (SMARTA TcR Transgenic), C57BL/6-Tg(Tcra2D2,Tcrb2D2)1Kuch/J (2D2 TcR Transgenic), B6.129S4-Ccr2tm1Ifc/J (CCR2 KO), and B6.Cg-Rag2tm1.1Cgn/J (RAG2 KO). Kaede transgenic mice were obtained from RIKEN. CAG-mOrange transgenic mice were provided by Dorian B. McGavern from NIH NINDS. Transgenice TCR transgenic mice were crossed CD45.1 congenic mice before being crossed to either Kaede, GFP, OFP, or PA-GFP mice. REX3 mice, which report the expression of CXCR3 ligands, were provided by Andrew D. Luster from Massachusetts General Hospital. REX3 mice were crossed with Albino B6 mice and subsequently with CCR2 KO or RAG KO mice.

### Method Details

#### T cell culture and adoptive transfers

For the *in vitro* generation of effector Th1 cells, naive CD4+ T cells were isolated from the lymph nodes and spleens of OT-II TCR transgenic mice using negative selection with a mouse Naive CD4+ T Cell Isolation Kit (Miltenyi Biotec) after treating with ACK lysis buffer to lyse red blood cell. Splenocytes from C57BL/6 mice were irradiated with 2500 rads to serve as antigen-presenting cells (APCs) after red blood cell lysis. To polarize cells to Th1 cells, naive CD4+ T cells were plated with the irradiated APCs at a ratio of 1:4, 1 μM cognate OVA peptide, IL-2 (10 U/ml), IL-12 (20 ng/ml; Peprotech) and anti-IL-4 (40 μg/ml; 11B11) in RPMI (Gibco) with supplemented FBS, 2-ME, L-Glutamine, Penicillin, and Streptomycin. Cells were incubated at 37°C with 5% CO2 and split at a 1:2 ratio three days later and harvested on day 5 or 6 of culture. For PA-GFP studies, cells were labeled with CellTracker DeepRed dye by incubating at 37°C in a 1:1000 working solution (Thermo Fisher Scientific). A total of 5×10^6^ OT-II TCR Th1 cells were adoptively transferred into recipient mice through retro-orbital injection on D2 post immunization. For co-transfer studies, 2.5×10^6^ each of OT-II Th1 and either 2D2 Th1 or SMARTA Th1 cells were transferred into recipient mice.

#### In vivo treatments and reagents

Mice were immunized intradermally (i.d.) in the ear with 10 μg OVA protein (Sigma Aldrich) emulsified in complete Freund’s adjuvant (CFA, Sigma Aldrich). For neutrophil depletion, rat IgG2a κ isotype control or anti-Ly6G antibody (200μg, 1A8, BioXCell) was administered intraperitoneally (i.p.) starting on day −1 prior to immunization and continued through day 4 post-immunization (p.i.) while co-administering mouse anti-rat Igk (400μg, MAR18.5, BioXCell) every other day^37^. CXCR2 antagonist SB225002 was administered i.p. at 50μg/200μL in PBS 0.5% Tween20 or with the vehicle PBS 0.5% Tween20 as a control.

#### Intravital multiphoton imaging

Mice were anesthetized with isoflurane (induction ∼4%; maintenance ∼1.5% with O_2_) using an isoflurane vaporizer-ventilation machine (Somnosuite, Kent Scientific).). For some experiments mice were anesthetized with 2,2,2-tribromomethanol (TBE, 240mg/kg) and maintained with isoflurane. Once anesthetized, the ventral side of the ear pinna was taped to a coverslip, and mice were placed on a custom-made platform for imaging^83^. Body temperature was maintained with a heated blanket (Kent Scientific) and a heating block (WPI). The microscope objective was heated to 36°C (Bioptechs) to maintain constant dermal temperature during imaging. Images were acquired with an Olympus FVMPE-RS twin-laser multiphoton system equipped with 4 photomultiplier tubes (PMTs) with blue, green, near-red, and far-red filters (Semrock). The FVMPE-RS platform utilizes the Olympus Fluoview software FV31S-SW software. Fluorescence was collected with an Olympus XLPlanN 25x objective (numerical aperture, 1.05) for deep-tissue multiphoton imaging and detected with three proprietary photomultipliers. Fluorescence excitation was achieved by a sequential scan of a Spectra-Physics DeepSee-MaiTai HP Ti laser tuned to 800 nm and a Spectra-Physics InsightX3 laser tuned to 920 nm with DM690-870 and DM690-1050 dichroic mirrors. The InsightX3 laser was tuned to 1080nm to visualize CCR2-mRFP when appropriate. The z-stack images (512 × 512 pixels) were 60 μm in thickness and acquired with a vertical resolution of 3 μm and a lateral pixel size of 994 nm. For larger-scale tiling of the ear the multi-area timelapse (MATL) function was used to record multiple single z-stacks in a registered area. Individual z-stacks were then made into a single cohesive image using the stich function in the Olympus Fluoview software. For time-series analyses, three-dimensional stacks were acquired approximately every 50 seconds. For acute detection of neutrophils in the inflamed ear, a 2:5:3 mixture of anti-Ly6G FITC (1A8, Biolegend) and anti-CD16/32 antibody (Fc Block, 2.4G2) was dialyzed against PBS to remove sodium azide. Immunized mice were injected intradermally in the ear with the antibody mix 2h prior to intravital multiphoton microscopy (IV-MPM) imaging. Blood and lymphatic vessels were stained with anti-CD31 AF647 (10 μg per mouse, administered retro-orbitally 15 minutes prior to imaging, clone 390, Biolegend). For Kaede photoconversion studies, on day 5 post-immunization (p.i.), the ear pinna was exposed to violet light^84^ for 10 minutes to ensure complete photoconversion from Kaede-green to Kaede-red of all cells in the ear. The rest of the mouse was shielded with aluminum foil, so cells in the blood and lymph nodes remained Kaede-green. Mice ear pinnae were imaged or harvested for flow cytometry immediately after photoconversion or 24h after exposure to violet light. For PA-GFP migration studies, mice ear pinnae were imaged on day 5 p.i., and BFP+ niches and DeepRed PA-GFP Th1 cells were identified by imaging at 800 nm, where photoactivation does not take place. Th1 cells were photoactivated to GFP+ using 50-70% laser power scanning at 830 nm ^85^. The photoactivation site coordinates were mapped on the Olympus user interface, and 24h after photoactivation, the original photoactivation site coordinates were identified on the ear pinnae and imaged.

#### Analysis of multiphoton data using Imaris and Python

Raw imaging data were processed with Imaris software (Bitplane). A 3×3 median filter was applied to reduce noise, and T cells were tracked using automated algorithms with manual correction. For co-transfer time-series, cell tracks lasting less than 8 minutes were excluded from analyses. No minimum cell-displacement criteria were imposed that would exclude non-motile cells. Average velocity was calculated in Imaris using the track speed mean. Arrest coefficients were determined as the ratio of the time a cell was not moving (instantaneous speed < 2 μm/min) to the total time the cell was observed. Cell tracks were linearly interpolated to maintain consistent time windows for analysis. Niche quantitation was performed using previously generated custom DBSCAN-based Python plugin^12^ for Imaris. Movies were created in Imaris, with animation and titles added using Adobe Premiere Pro.

#### Relative cell distance

PA-GFP^+^ T cells and CXCL10^+^ Cells were volumetrically rendered in 3D using Imaris and Niches identified using DBSCAN-based python plugin. Imaris Distance Transformation Xtension was applied to the PAGFP^+^ or CXCL10^+^ surface, resulted in a new channel with the intensities equal to the distance from the surface object. Minimal distance of each PAGFP^+^ T cell from the CXCL10^+^ niche was calculated using Imaris.

#### Isolation of CXCL10+ Enriched Niche and Non-Niche Regions

Prior to mounting the ear pinna to its coverslip as described above, two small marks were made with a marker on the superior and lateral edges of the inside ear. After mounting, the (x, y) coordinates of these two marks were registered in the Olympus software’s map feature. The ear was then manually scanned and any visible clusters of BFP+ cells were marked on the coordinate map, and adjacent regions devoid of BFP signal were noted as well. Following this the ear was removed from the coverslip and placed on a flat surface. Then using a 100μm scale ruler, the niche regions identified (along with adjacent non-niche) were located on the x, y plane in relation to the two marker points. After confirming the location of the niches in relation to both points, a 2mm bore biopsy punch was centered over each coordinate and used to isolate niche and non-niche and niche enriched regions. Two niche and two non-niche biopsies were taken per ear to control across animals and experiments, selecting for regions containing the most BFP signal for the niche sample in the instance more than two were located.

#### Flow Cytometry

Mice were euthanized, and their ears were excised. For experiments with the adoptive transfer of Kaede cells, mice were given 3 ug of CD45 BV650 one minute prior to euthanizing. Ears was then split into ventral and dorsal sheets and placed in 1.5 ml of Liberase TL (0.4 mg/mL in RPMI; Sigma Aldrich) and 50ug of DNase I (Sigma Aldrich) per ear. Ears were minced and incubated with agitation for 1h at 37°C. Tissue digestion was quenched by adding complete media (RPMI 1640 supplemented with 10% fetal calf serum, Lonza), followed by mechanical disaggregation using program C on the GentleMACS dissociator (Miltenyi Biotec). Insoluble material was removed by filtration, and the pellet was washed twice with 2% neonatal calf serum in Hank’s balanced-salt solution (HBSS). Single cell suspensions were stained with Ghost Dye UV 450 or Violet 510 dead cell stain for 405 nm excitation (Tonbo Biosciences) at a 1:1000 dilution in PBS for 30 minutes at 4°C, followed by a wash with 2% NCS in HBSS. Cells were then incubated for 10 minutes with a 1:100 dilution of CD16/CD32 Fc block (2.4G2) in FACS buffer (2% NCS/PBS), followed by staining for 30 minutes at 4°C with a fluorochrome-conjugated antibody mix in FACS buffer. Antibodies used include: anti-CD45 BUV395 (1:500, 30-F11, BD Biosciences), anti-CD4 BUV805 (1:500, GK1.5, BD Biosciences), anti-PD1 BV605 (1:100, J43, BD Biosciences), anti-CD137 PE-Cy7 (1:100, 17B5, Invitrogen), anti-CD153 APC (1:100, RM153, Invitrogen), anti-PD1 PE-Cy7 (1:100, J43, Invitrogen), anti-CD137 APC (1:100 CD137, 17B5, Invitrogen), anti-CD64 BV605 (1:200, X54-5/7.1, Biolegend), anti-CD11c BV711 (1:450, N418, Biolegend), anti-CD19 BV786 (1:100, 1D3, BD Biosciences), anti-F4/80 PE-Cy5 (1:200, BM8, Invitrogen), anti-CD88 anti-APC (1:800, 20/70, Biolegend), anti-MHC-II APC-ef780 (1:2000, M5/114.15.2, eBioscience), anti-CD26 PE-Cy7 (1:50, H194-112, Biolegend), anti-XCR1 BV650 (1:200, ZET, Biolegend), anti-Ly6G FITC (1:500, 1A8, Biolegend), anti-CD3e Biotin (1:400, 145-2C11, eBioscience), anti-CD4 BV605 (1:200, RM4-5, Biolegend), anti-CD4 (1:200, RM4-4, Biolegend), anti-CD45.1 BUV563 (1:250, A20, BDbiosciences), and anti-CXCR6 Biotin (1:100, SA051D1, Biolegend). Cells stained with biotinylated antibodies were then washed and incubated with either Streptavidin BV510 (1:1000, Biolegend) or Streptavidin PE-Cy7 (1:500, Biolegend) diluted in FACS buffer for 30 minutes at 4°C. The cells were washed, filtered, and resuspended in FACS buffer containing 100μg/mL of DNase I and 20 μL of AccuCheck counting beads (Thermo Fisher Scientific) per ear. Samples were collected using a FACSymphony A3 (BD Biosciences) and analyzed with FlowJo software (Treestar, Ashland, OR).

#### Luminex

Mice ears were excised, digested, and disaggregated as described previously, in a total volume of 500 μl, and then kept on ice. The homogenates were centrifuged at 13,000rpm, and the supernatant was collected and stored at −80°C until analysis where samples were passed through a 40μm filter and re-centrifugated prior to use. 50μl of neat or 1:10 diluted ear homogenate was assayed for chemokines and cytokines using the Milliplex MAP Mouse Cytokine/Chemokine Magnetic Bead Panel kits I & II (Millipore) the Samples were collected and analyzed with the MAGPIX System (Diasorin).

#### ELISpot

Single cell suspensions of ears were generated as described above and from lymph node by gently reducing lymph nodes over 70μm cell filters. Single cells were suspended in RPMI (Gibco) supplemented with FBS, 2-ME, L-Glutamine, Penicillin, and Streptomycin for plating. ELISpot plates (MilliPore, ref: MSIPN4W50) were prepared by coated with anti-mouse IL-2 (1:250, JES6-1A12, BD Pharmagen) or IFNγ (1:250, AN18, made in house) O/N at 4°C. Plates were then blocked with 10% FBS containing media for 2h at RT. Serial dilutions of cells were plated in coated and blocked plates. Cells were serially diluted in duplicate with wells receiving either no Ag or 1µM pOVA for restimulation. Cells were cultured at 37°C for 18h with 5% CO2. Biotin anti-mouse IL-2 (1:250, JES6-5H4, BD Pharmagen) or IFNγ (1:250, XMG, eBioscience) Ab was added to wells for 1h at RT. Streptavidin alkaline phosphatase (1:1000, Jackson Laboratories) was then added to wells for 30 minutes at RT followed by development with Vector Blue Alkaline Phosphatase Substrate Kit III (Vector Laboratories, SK-5300). After drying plates were read using the ImmunoSpot platform (Cellular Technology Limited).

#### Single-cell RNA sequencing

Single-cell suspensions were submitted to the Cornell BRC Genomics Facility for processing and sequencing. The Chromium Next GEM Single Cell 3’ reagent kit v3 from 10X Genomics® protocol was used for FACS-sorted cell sequencing. A target of 10000 cells across runs were loaded for Gel Beads-in-emulsion (GEM) generation. According to the kit protocol first GEMs were generated, subsequently reverse transcription was performed, and cDNA was cleaned up, amplified, and size selected. Sample quality was confirmed using a Qubit (RNA HS kit; Thermo Fisher) to determine concentration and a Fragment Analyzer (Agilent) to determine RNA integrity. The libraries were sequenced with an Illumina NextSeq 2000.

#### Single-cell RNA-seq: data processing

Raw sequencing reads for the mRNA libraries were processed into raw count matrices using CellRanger version 6.1.2 (10X Genomics) with the mm10 2020-A reference. The raw counts for mRNA were analyzed in R through the RStudio integrated development environment using Seurat v4.1.2 ^86^. Data were filtered to include only single cells with between 200 and 5000 genes and less than 15% mitochondrial reads. Normalization was carried out using sctransform, followed by identification of variable features and scaling of the data. Samples were integrated using Harmony ^87^, and clustering was performed in Seurat by using 12 dimensions with the FindNeighbors function and a resolution of 0.5 for the FindClusters function. Marker genes for each cluster were identified using Seurat’s FindMarkers function. The Tidyverse, dittoSeq, ComplexHeatmap, and various Seurat functions were utilized for plotting the scRNA-seq data.

#### Pseudobulk Analysis

Differential expression analysis between Niche and Non-Niche samples was performed using the FindMarkers function with MAST testing^88^ to identify the top genes expressed in each condition. Clusters were subsetted from the original Seurat object, and log_2_-fold change matrices were generated through comparisons within the Niche and Non-Niche cells of each cluster. P-values from the log_2_-fold change matrices were adjusted using the Benjamini & Hochberg method, with results filtered for adjusted p-values below 0.05 and sorted by log2-fold change to establish a final gene rank list. Gene set enrichment analysis was conducted using clusterProfiler v4.0^89^, with Gene Ontology domains of Biological Process and Molecular Function. Myeloid cell and T-cell signatures were derived from a unique list of genes derived from the GSEA leading edge genes. Signature scores were calculated per sample with the R package UCell v2.20^90^.

#### Ligand-Receptor Pair Analysis

CellChat v2.1.0^91^ was utilized to infer cell-cell communication probabilities and identify signaling changes across cell populations. The Secreted Signaling mouse CellChat database was used for cell-cell communication analysis. Differentially overexpressed ligands and receptors were identified, and their average expression values were used to calculate communication probabilities among all cell clusters. Interaction strength along specific intercellular signaling pathways was assessed by aggregating the communication probabilities of associated ligand-receptor pairs across clusters. To assess the contribution of cells expressing ligands or receptors within the CCL and CXCL communication pathways, communication probabilities for cells that act as senders (ligand-expressing) and receivers (receptor-expressing) were calculated.

#### Quantification and Statistical Analysis

Statistical analyses were performed using GraphPad Prism or R software. Nonparametric Mann-Whitney and Kolmogorov-Smirnov tests were primarily utilized to compare two treatment groups. For analyses involving multiple comparisons, the Kruskal-Wallis test followed by Dunn’s post-test was employed. Analysis of variance (ANOVA) was used to assess differences across multiple groups. Significance levels were denoted as follows: *p ≤ 0.05, **p ≤ 0.01, ***p ≤ 0.001, ****p < 0.0001, and N.S. (not significant) for p > 0.05.

