## Supplementary material for "Th1 cells are critical tissue organizers of myeloid-rich perivascular activation niches": Bala, McGurk et al Supplementary Figures

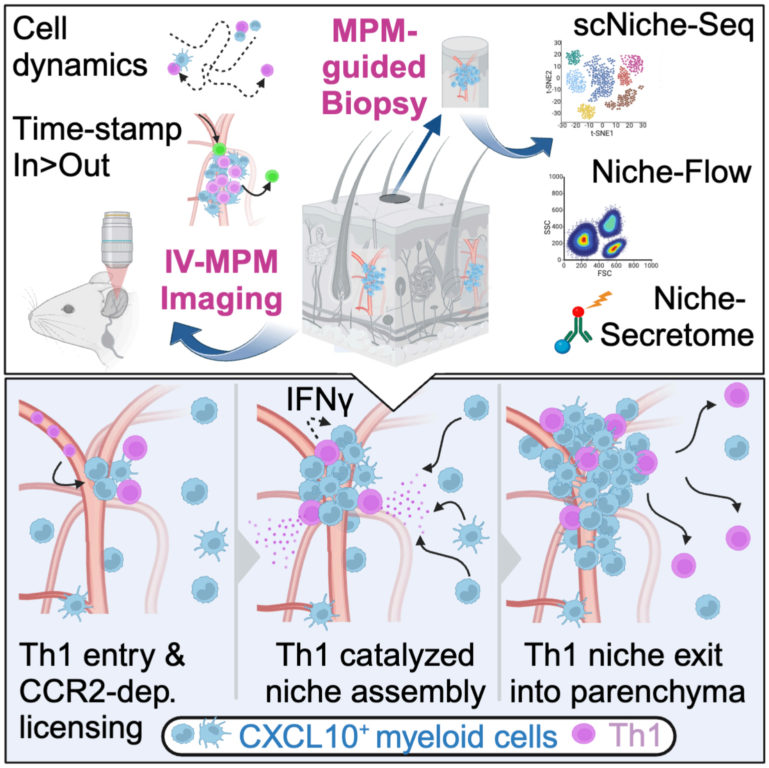
Cell
dynamics
Time-stamp
In>Out
MPM-
guided
Biopsy
scNiche-Seq
Niche-Flow
IV-MPM
Imaging
Niche-
Secretome
IFNγ
Th1 entry &
CCR2-dep.
licensing
Th1 catalyzed
niche assembly
Th1 niche exit
into parenchyma
CXCL10+ myeloid cells
Th1

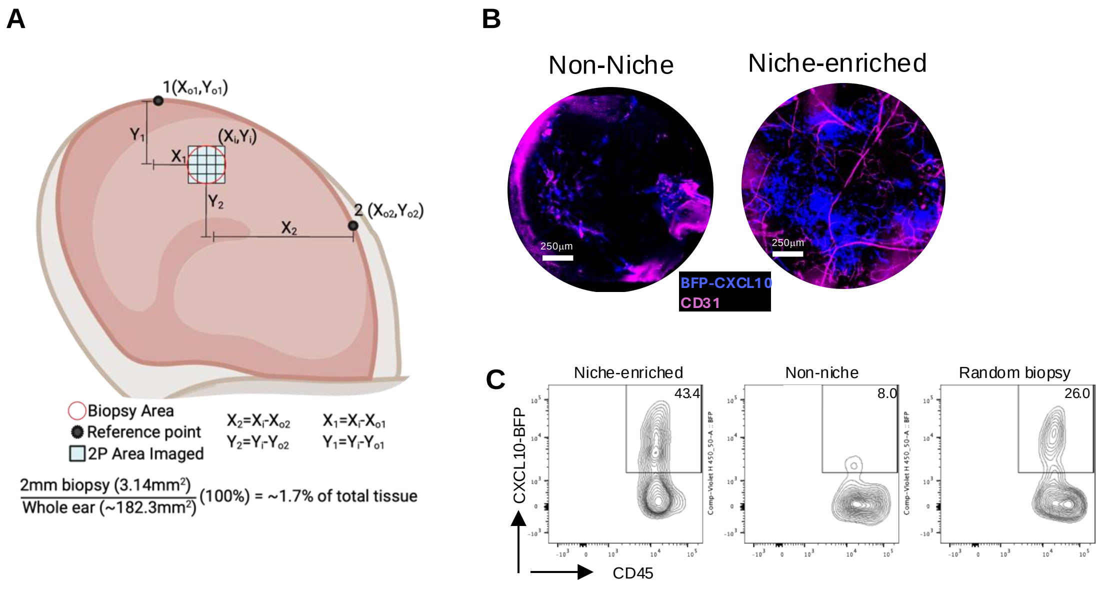


**Figure S1. Multiphoton microscopy-guided identification of niche regions for biopsy.** REX3 mice were immunized with OVA/CFA and IV-MPM preformed d5 post-immunization, IV-labeling of CD31 with Alexa 647 (pink) 30 mins prior to imaging. A) Spatial coordinates of reference points made on the outer edge of the ear pinna with permanent marker, as well as CXCL10-BFP PVN rich regions (niche-enriched) and CXCL10-BFP devoid regions (non-niche) by MPM were mapped with the Olympus user interface "register area" function. Using coordinates from each reference point and individual registered area, x (um) and y (um) distances were calculated and checked for agreement between reference points 1 and 2. Regions identified by these calculated distances were then biopsied (2 mm punch biopsy) and subsequently reimaged. B) Representative image of niche-enriched and non-niche 2 mm punch-biopsies. C) Flow cytometry of MPM guided biopsy material from niche-enriched and non-niche regions of the inflamed ear pinna, compared to random punch biopsies of the inflamed ear pinna.

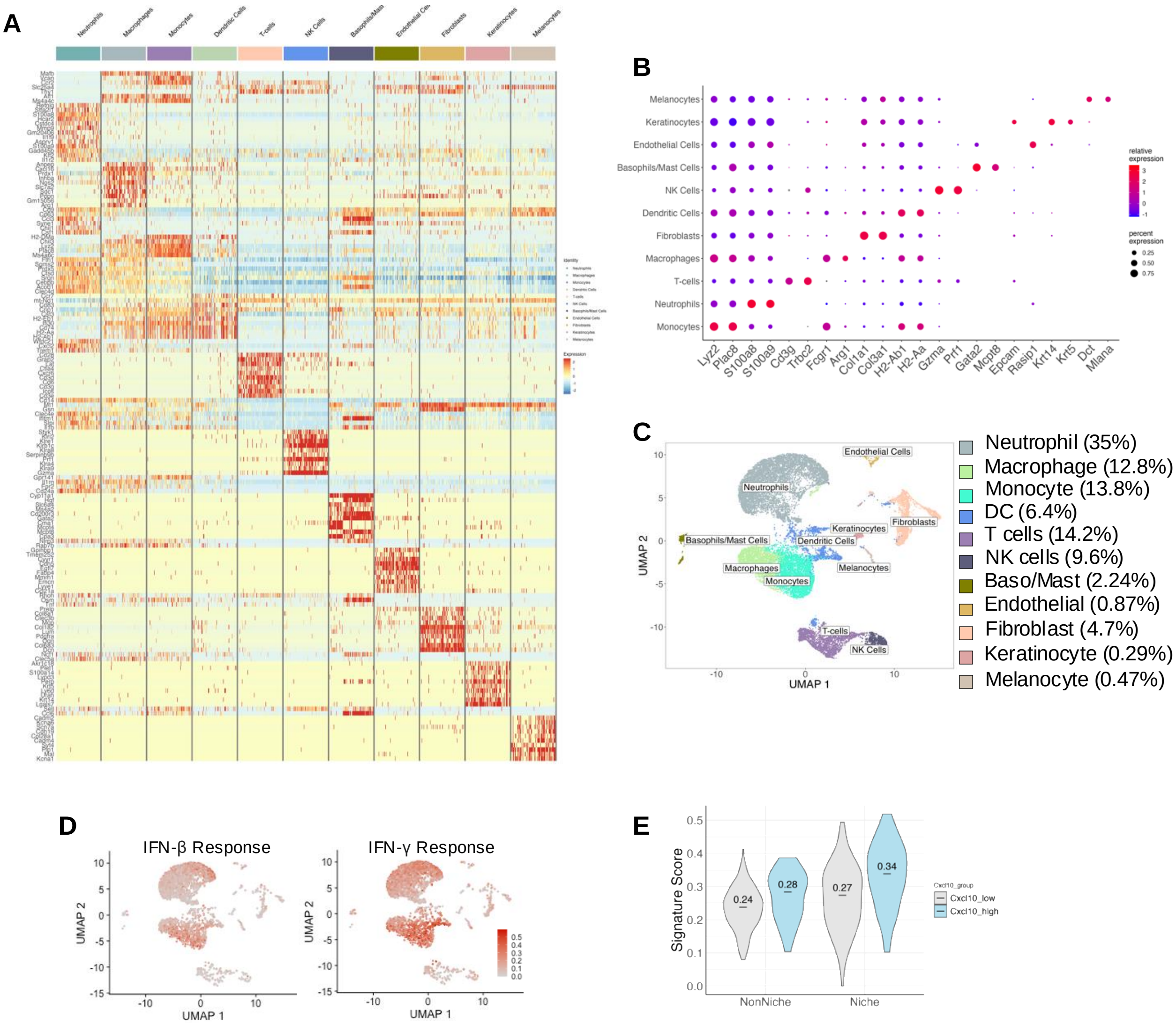


**Figure S2. Spatially-resolved scRNAseq cell clustering and analysis.**
REX3 mice were immunized with OVA/CFA and IV-MPM preformed d5 post-immunization, with IV-labeling of CD31 with Alexa 647 (pink) 30 mins prior to imaging. Niche and non-niche biopsies, as well as whole tissue (ear pinna) were processed for scRNAseq. (A) Heatmap of top 10 genes represented in each cell cluster from UMAP representation. FindMarkers function in Seurat was used to determine marker genes for distinguishing each cluster from the remaining cluster. Row Z-score represents normalized expression across each gene. (B) Dotplot of two identifier genes per cluster used for annotation of each cluster identified within the UMAP. (C) Quantified cluster distribution percentages within the whole tissue. (D) Niche scored by responsiveness to type I IFN (IFN- β) or IFNγ and overlaid on UMAP. (E) IFNγ signature score in CXCL10$^{hi}$ and CXCL10$^{lo}$ monocytes in niche and non-niche regions.

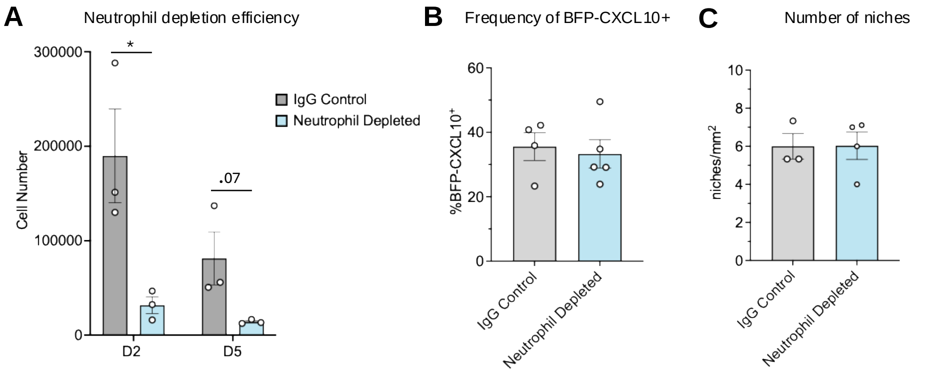


**Figure S3. Sustain neutrophil recruitment is not required for CXCL10 induction or niche formation.** (A) Quantification of neutrophil infiltrate in d2 and d5 CFA/OVA immunized ears following treatment with 200mg non-specific IgG isotype control mAb or anti-Ly6G mAb (1A8) and 400 mg anti-mouse kappa light chain mAb (MAR 18.5). Neutrophils were pre-gated on Live/CD45$^+$/CD4$^-$/CD64$^-$/F4/80$^-$/MHCII$^-$ cells and identified as CD11b$^+$ Ly6C$^+$ cells, excluding Ly6C$^{hi}$ monocytes. (B) Frequency of BFP-CXCL10$^+$ cells in d5 CFA/OVA immunized ears, pre-gated on Live/CD45$^+$ cells. (C) Quantification of niche formation in d5 CFA/OVA immunized ears per mm$^2$. (A-C) Statistical analysis performed using Mann-Whitney test; *$p<0.05$

| PVN Gene Signature | |
|---|---|
| CCL4 | B2M |
| CCL3 | PSMB6 |
| CXCL9 | HLA-DRA |
| CXCL10 | HLA-DRB1 |
| TLR2 | HLA-DQB1 |
| NFKBIA | CIITA |
| IRF1 | CD80 |
| CD14 | CD86 |
| TNF | GBP3 |
| CSF2RB | IRGM |
| PRDX5 | GBP6 |
| GLRX | GBP5 |
| PRDX1 | TAP1 |
| TXNRD1 | GBP7 |
| SOD2 | IFI16 |
| SLPI | NLRC5 |
| TIMP1 | HLA-B |
| IFITM1 | HLA-G |
| PTGS2 | GBP4 |
| HLA-C | PSMB9 |
| | TIMP1 |

**Figure S4. PVN myeloid gene signature.** The PVN gene signature was compiled with genes enriched in the niche in 3 out of 4 myeloid cell populations: neutrophils, monocytes, macrophages and dendritic cells. In addition, niche-enriched genes associated with IFN$\gamma$ and antigen presentation that were elevated in PVN monocytes.

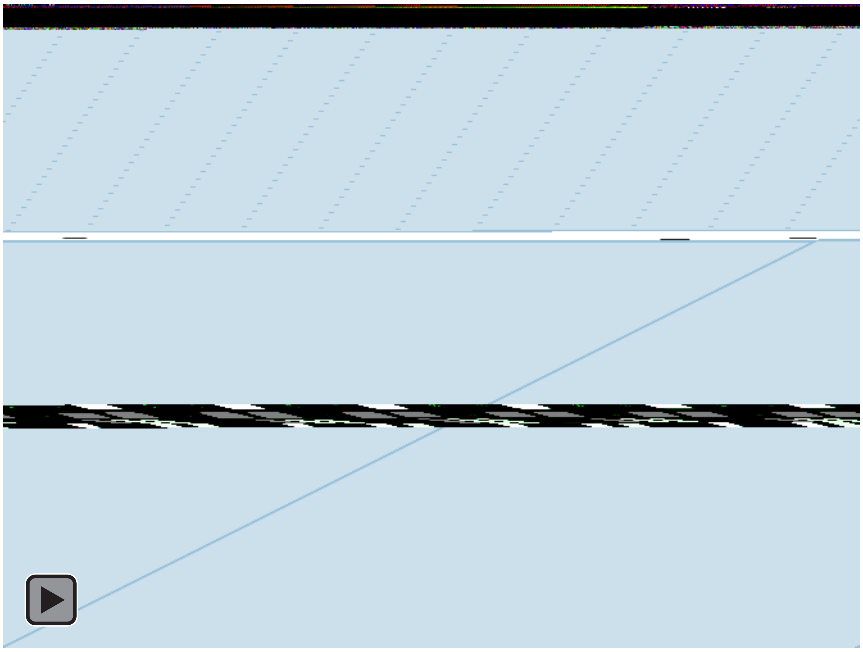

**Video S1. Antigen-specific Th1 arrest in the perivascular niche.**
*In vitro* generated LCMV $GP_{61\text{-}80}$-specific (SMARTA) Th1 OFP cells were intravenously co-transferred alongside ovalbumin-specific (OT-II) Th1 GFP cells into REX3 mice intradermally immunized with OVA/CFA in the ear pinna. Day 5 post-immunization IV-MPM was used to visualize the inflamed dermis and identify the CXCL10-BFP$^{+}$ PVN. Distribution of BFP-CXCL10$^{+}$ cells (blue) in the dermis and motility tracks of OT-II GFP$^{+}$ (green) and SMARTA OFP$^{+}$ (red). The video represents a 2D maximal z-projection time series of a 60µm-thick imaging volume; scale bar 80µm.
